# Spatial tuning shifts increase the discriminability and fidelity of population codes in visual cortex

**DOI:** 10.1101/086892

**Authors:** Vy A. Vo, Thomas C. Sprague, John T. Serences

**Affiliations:** Neurosciences Graduate Program, University of California, San Diego, La Jolla, CA 92093; Department of Psychology and Center for Neural Science, New York University, New York, New York 10003; Department of Psychology, University of California, San Diego, La Jolla, CA 92093; Kavli Institute for Brain and Mind, University of California, San Diego, La Jolla, CA 92093

## Abstract

**ABSTRACT:** Selective visual attention enables organisms to enhance the representation of behaviorally relevant stimuli by altering the encoding properties of single receptive fields (RFs). Yet we know little about how the attentional modulations of single RFs contribute to the encoding of an entire visual scene. Addressing this issue requires (1) measuring a group of RFs that tile a continuous portion of visual space, (2) constructing a population-level measurement of spatial representations based on these RFs, and (3) linking how different types of RF attentional modulations change the population-level representation. To accomplish these aims, we used fMRI to characterize the responses of thousands of voxels in retinotopically organized human cortex. First, we found that the response modulations of voxel RFs (vRFs) depend on the spatial relationship between the RF center and the visual location of the attended target. Second, we used two analyses to assess the spatial encoding quality of a population of voxels. We found that attention increased fine spatial discriminability and representational fidelity near the attended target. Third, we linked these findings by manipulating the observed vRF attentional modulations and recomputing our measures of the fidelity of population codes. Surprisingly, we discovered that attentional enhancements of population-level representations largely depend on position shifts of vRFs, rather than changes in size or gain. Our data suggest that position shifts of single RFs are a principal mechanism by which attention enhances population-level representations in visual cortex.

**SIGNIFICANCE STATEMENT:** While changes in the gain and size of RFs have dominated our view of how attention modulates information codes of visual space, such hypotheses have largely relied on the extrapolation of single-cell responses to population responses. Here we use fMRI to relate changes in single voxel receptive fields (vRFs) to changes in the precision of representations based on larger populations of voxels. We find that vRF position shifts contribute more to population-level enhancements of visual information than changes in vRF size or gain. This finding suggests that position shifts are a principal mechanism by which spatial attention enhances population codes for relevant visual information in sensory cortex. This poses challenges for labeled line theories of information processing, suggesting that downstream regions likely rely on distributed inputs rather than single neuron-to-neuron mappings.

## INTRODUCTION

Spatial receptive fields (RFs) are a core component of visual information processing throughout the visual system. They are modified by selective visual attention to improve the fidelity of sensory representations, likely enabling more precise, accurate perception (Desimone and Duncan, 1995; Anton-Erxleben and Carrasco, 2013). Prior studies in non-human primates have found that covert spatial attention changes the position, size, and amplitude of responses in single-cell RFs in early cortical areas such as V1, V4, and MT (Moran and Desimone, 1985; Connor et al., 1996, 1997, Womelsdorf et al., 2006, 2008; Roberts et al., 2007; David et al., 2008). Recent neuroimaging studies have also shown that single-voxel RFs (vRFs) undergo similar response changes with attention, shifting towards the attended target or changing in size (de Haas et al., 2014; Klein et al., 2014; Kay et al., 2015; Sheremata and Silver, 2015). Most accounts suggest that these RF modulations improve the spatial representations of the attended target, either by boosting the signal-to-noise ratio (SNR) by increasing response amplitude, or by increasing the spatial resolution by decreasing RF size (Desimone and Duncan, 1995; Anton-Erxleben and Carrasco, 2013; Cohen and Maunsell, 2014). These mechanisms are akin to turning up the volume (gain increase) or to using smaller pixels to encode a digital image (size decrease).

Despite these documented modulations, it is not yet clear how different types of RF modulations are combined to facilitate robust population codes. Recent studies have only begun to explore how interactions between neurons may affect the coding properties of the population (Anton-Erxleben and Carrasco, 2013; Cohen and Maunsell, 2014). Yet analyzing these data at a population level is crucial for understanding how spatial attention changes the overall representation of an attended area. Prior fMRI studies that measured many vRFs across space were often unable to report the full pattern of response modulations with respect to the attended target because subjects attended to the mapping stimulus, rather than to a fixed point in space (Sprague and Serences, 2013; Kay et al., 2015; Sheremata and Silver, 2015). Studies which fixed the locus of attention have reported mixed results on vRF modulations (de Haas et al., 2014; Klein et al., 2014). The first aim of this study was thus to evaluate how properties of vRFs in retinotopic areas change with attention, especially near the peripheral attention target.

The second aim of the study was to evaluate how different types of RF modulations contribute to population-level enhancements of an attended region of space. Single RFs in early visual areas are fundamentally local encoding models that are relatively uninformative about regions outside their immediate borders. To study their relationship to a population-level representation of space, other metrics are needed to integrate information across all local encoding units – e.g., vRFs – to evaluate how attentional modulations impact the quality of population codes. Here, we used two different population-level metrics of spatial encoding fidelity to investigate these questions, and to determine how changes in vRF amplitude, size, or position affect the population-level representations. First, we used a measure related to Fisher Information to evaluate the spatial discriminability of population codes. Second, we used a spatial encoding model that incorporates information across voxels to form representations of stimuli in the mapped visual field (Brouwer and Heeger, 2009; Sprague and Serences, 2013; Sprague et al., 2015).

We found that vRF position shifts increase both the spatial discriminability around the attended region as well as the fidelity of stimulus reconstructions near the attended target. Surprisingly, shifts in vRF position accounted for more of the population-level enhancements with attention than changes in vRF size or gain. This finding is unexpected in the context of ‘labeled-line’ models of information processing, which posit that visual representations rely on RFs that transmit consistent ‘labels’ for visual features such as spatial position. Our findings suggest that apparent shifts in the labels of RFs play an important role in the attentional enhancement of visual information.

## MATERIALS & METHODS

### Task design and participants

We collected data from 9 human participants (4 female), 6 of whom had previously completed a set of retinotopic mapping scans in the lab (participants AA, AB, AC, AI, and AL in Sprague & Serences, 2013; participants AA, AC, and AI in Sprague et al., 2014; all participants in Ester et al., 2015). All participants provided written informed consent and were compensated for their time ($20/hour) as approved by the local UC San Diego Institutional Review Board. Participants practiced both the attention task and the localizer task before entering the scanner. A minimum of four hours of scanning was required to complete the entire analysis, so one participant was excluded due to insufficient data (they only completed 2 hours). Another participant was excluded for inconsistent behavioral performance, with average task accuracy at chance (48.6%). This yielded a total of 7 participants who completed the entire experiment (3 2-hour scan sessions per participant).

Participants centrally fixated a gray rectangular screen (120 × 90 cm) viewed via a head-coil mounted mirror (~3.85 m viewing distance). They attended one of three fixed locations on the screen: the fixation point or a target to the lower left or lower right of fixation. During each 2000 ms trial, subjects reported a change in the attention target. When subjects attended fixation, they reported whether a brief contrast change (100 – 400 ms, starting 300 – 1000 ms into the trial) was dimmer or brighter than the baseline contrast. The peripheral attention targets were two pentagons (0.17° radius; 50% contrast) centered 2.1º to the left and right of fixation (Fig 1a). When subjects attended a peripheral target, they reported whether it rotated clockwise or counter-clockwise (rotation duration 100 - 300 ms, starting 300 - 1600 ms into the trial). Inter trial intervals (ITIs) randomly varied between 1000 to 3000 ms in 500 ms increments (mean ITI: 2000 ms). The magnitude of the contrast change or the rotation was adjusted on each run to keep task performance for each participant near 75% (mean = 75.90%, bootstrapped 95% C.I. [72.46%, 79.20%]), with no significant difference between conditions as evaluated with a one-way repeated measures ANOVA randomization test (F(1,11) = 0.220, randomized p = 0.800). For four participants, we collected 6 runs on the attend periphery tasks without a change in the luminance of the fixation stimulus. Performance on the attend periphery tasks was stable across runs with and without the luminance change (repeated-measures ANOVA with run type x random participants factor; p = 0.439, null F distribution using randomized labels for 10,000 iterations). Therefore, these data were collapsed across scan sessions with and without changes in fixation luminance.

**Figure 1.**
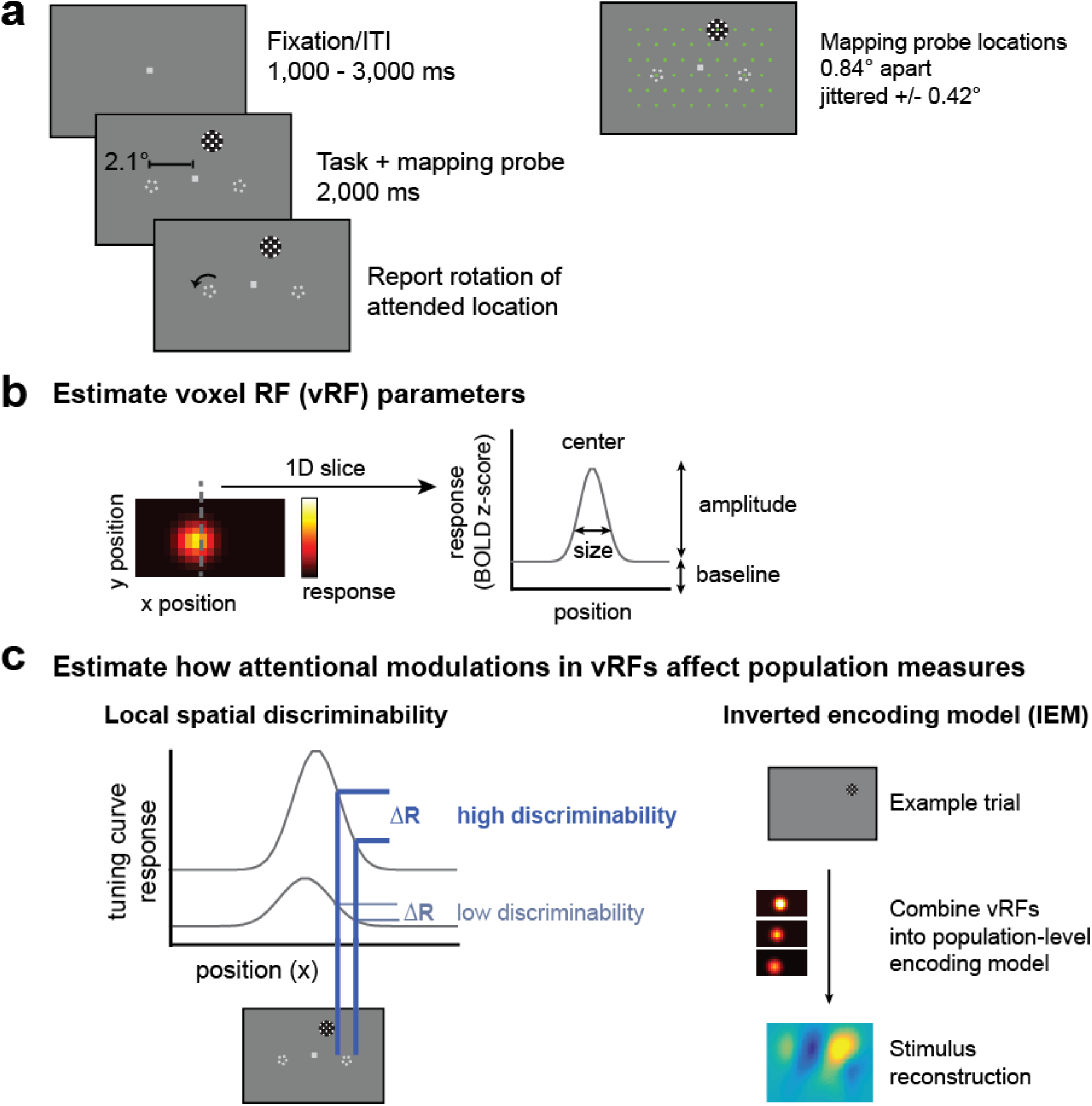
Covert spatial attention task and hypothesized representation changes with shifts of spatial attention. (**a**) Subjects fixated centrally and attended to brief rotations in the pentagon stimulus on the left or right while a flickering checkerboard probe stimulus appeared at one of 51 grid locations across the visual field. On control runs, subjects attended to a contrast change at fixation. fMRI data measured during this attention task is used to create visualizable estimates of voxel receptive fields (vRFs) and stimulus reconstructions. (**b**) A receptive field model is fit to the responses of each voxel, and can be described by its x and y position (center), its response baseline, response amplitude, and its size (full-width half maximum). (**c**) Given a population of voxels in a retinotopic region, such as V1, we examine two different measures of spatial information in the population. The first, a spatial discriminability metric, scales with the slope of the tuning curve at a given location in space (**Materials and Methods**). The second relies on a multivariate inverted encoding model (IEM) for space. By reconstructing images of the mapping stimulus on each test trial, we can measure how population-level spatial information changes with attention. We then can model how changes in individual vRFs affect both of these population measures.

On 51 of the 61 trials in each run, a full-contrast 6 Hz flickering checkerboard (0.68° radius; 1.67 cycles/deg) appeared for 2000 ms at one of 51 different locations across the screen to map the spatial sensitivity of visually responsive voxels. These mapping stimuli covered a region of the screen roughly subtending 9° horizontal and 6° vertical when their position was jittered. When one of these checkerboards overlapped with any of the static attention targets, they were partially masked with a small circular aperture the same color as the screen background (0.16°/0.25° radius aperture for fixation/pentagon, respectively) that allowed the stimulus to remain visible. Participants were instructed to ignore the task-irrelevant flickering checkerboards throughout the experiment. During the 10 null trials on each scan, the participant continued to perform the attention task but no checkerboard was presented. Null trials and mapping stimulus trials were presented in a pseudorandom interleaved order.

The location of the checkerboard mapping stimulus on each trial was determined by generating an evenly spaced triangular grid (0.84° between grid points) and centering the checkerboard on one of these grid points. The location of the checkerboard mapping stimulus was then jittered a random amount from these grid points (+/− 0.42°/0.37° horizontal/vertical). When subjects attended the peripheral target, half of the runs were presented at the discrete grid positions so that we could achieve more stable stimulus reconstructions (see **Population analysis (2)**).

### Magnetic resonance imaging

We obtained all structural and functional MR images using a GE 3T MR750 scanner at University of California, San Diego. We collected all functional images (19.2 cm × 19.2 cm FOV, 64 × 64 acquisition matrix, 35 interleaved slices, 3 mm^3^ voxels with 0 mm slice gap, 128 volumes per scan run) using a gradient echo planar pulse sequence (2000 ms TR, 30 ms TE, 90° flip angle) and a 32-channel head coil (Nova Medical, Wilmington, MA). Five dummy scans preceded each functional run. A high-resolution structural image was acquired at the end of each session using a FSPGR T1-weighted pulse sequence (25.6 cm × 25.6 cm FOV, 256×192 acquisition matrix, 8.136/3.172 ms TR/TE, 192 slices, 9° flip angle, 1 mm^3^ voxels). All functional scans were co-registered to the anatomical images acquired during the same session, and this anatomical was in turn co-registered to the anatomical acquired during the retinotopy scan.

EPI images were unwarped with a custom script from UCSD’s Center for Functional Magnetic Resonance Imaging using FSL and AFNI. All subsequent preprocessing was performed in BrainVoyager 2.6.1, including slice-time correction, six-parameter affine motion correction, and temporal high-pass filtering to remove slow signal drifts over the course of each run. Data were then transformed into Talairach space and resampled to have a 3×3×3 mm voxel size. Finally, the BOLD signal in each voxel was transformed into Z-scores on a scan-by-scan basis. All subsequent analyses were performed in MATLAB using custom scripts (available online on Open Science Framework: osf.io/s9vqv).

### Independent localizer task

We constrained our analyses to visually responsive voxels in occipital and parietal cortex using a separate localizer task (3-5 runs per participant). On 14 trials, participants fixated centrally and viewed a full-field flickering checkerboard (10 Hz, 11.0/8.3° width/height) for 8000 ms. Participants detected whether a small area (2D Gaussian, σ = 0.2°) within the checkerboard dimmed in contrast. Contrast dimming occurred between 500 to 4000 ms after the start of the trial, and lasted between 2000 to 3000 ms (all uniformly sampled in 500 ms steps). This contrast change occurred infrequently (randomly on 5 out of 14 trials) at a random location within the checkerboard. The average contrast change was varied between runs to maintain consistent performance at ~75% accuracy (mean performance 78.0%). On 8 trials participants simply fixated throughout the trial without a checkerboard being presented. We then used a standard general linear model (GLM) with a canonical two-gamma hemodynamic response function (HRF, peak at 5 s, undershoot peak at 15 s, response undershoot ratio 6, response dispersion 1, undershoot dispersion 1) to estimate the response to the checkerboard stimulus in each voxel. For all subsequent analyses, only voxels in the retinotopically defined areas V1, V2, V3, V4, V3A/B and IPS0 with a significantly positive BOLD response to the localizer task (at FDR q = 0.05) were included (Benjamini and Yekutieli, 2001).

### Estimating single trial BOLD responses

For all subsequent analyses, we used trial-wise BOLD z-scores. We estimated these by creating a boxcar model marking the duration of each checkerboard mapping stimulus and convolving it with a canonical two-gamma HRF (peak at 5 s, undershoot peak at 15 s, response undershoot ratio 6, response dispersion 1, undershoot dispersion 1). To standardize our data across runs, we z-scored the BOLD responses within each run and concatenated the z-scores across runs. We then solved a GLM to find the response to each predictor.

### Statistical procedures

All reported confidence intervals (CIs) are computed by resampling the data with replacement (i.e. bootstrapping). The number of iterations for each bootstrapping procedure varied (depending on available computing power and time for that procedure) and are therefore reported with each result. For tests comparing a bootstrapped distribution against zero, p-values were computed by conducting two one-tailed tests against 0 (e.g., mean(param_change < 0) & mean(param_change > 0)) and doubling the smaller p-value. All repeated tests were FDR corrected (q = 0.05).

### Voxel receptive field (vRF) estimation, fitting, and parameter analysis

We first estimated vRFs for each attention condition to investigate (1) how vRF parameters changed when participants attended to different locations and (2) the spatial pattern of vRF changes across visual space. We note here that prior reports have referred to similar voxel RF models as population receptive fields, or pRFs, to emphasize the fact that each voxel contains a population of spatially tuned neurons (Dumoulin and Wandell, 2008; Wandell and Winawer, 2015). However, since we are comparing modulations at different scales in the present study (i.e. modulations in single voxels and in patterns of responses across many voxels), we will refer to these single voxel measurements as voxel receptive fields (vRFs), and will reserve the term ‘population’ exclusively for multivariate measures involving several voxels, allowing our terminology to align with theories of population coding (Ma et al., 2006).

We estimated voxel receptive fields (vRFs) using a modified version of a previously described technique (Sprague and Serences, 2013). This method estimates a single voxel’s spatial sensitivity by modeling its BOLD responses as a linear combination of discrete, smooth spatial filters tiled evenly across the mapped portion of the visual field. These spatial filters (or spatial channels) form our modeled basis set. We then regressed the BOLD z-scores (*v* voxels x *n* trials) onto a design matrix with predicted channel responses for each trial (*C*, *k* channels x *n* trials) by solving Equation 1:

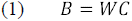

for the matrix *W*(*v* voxels x *k* channels).

Each of the *k* channels in the basis set was defined as a two-dimensional cosine that was fixed to reach 0 at a set distance from the filter center:

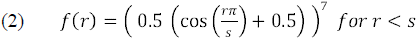

where *r* is the distance from the filter center and *s* is the size constant. Setting a zero baseline in this function ensured that we could estimate a stable baseline for each voxel by restricting the response of the channel to a known subregion of the visual display. Since the estimated vRF size depends on the size of the filters, we made the filters fairly small (1.08° FWHM) and dense (91 filters arranged in a 13 horizontal / 7 vertical grid, each spaced 0.83° apart). We then discretized the filters by sampling them in a high-resolution 2D grid of 135 by 101 pixels spanning 10° by 5°. The discretized filters (*k* filters by *p* pixels) were multiplied with a mask of the checkerboard stimulus on every trial (*p* pixels by *n* trials) so that the design matrix *C* contained predictions of the spatial channel responses on every trial of the mapping task.

To fit our estimated vRFs with a unimodal function, we used ridge regression to solve Equation 1. This is a common regularization method which sparsifies the regression solution by penalizing the regressors with many small weights (Hoerl and Kennard, 1970; Lee et al., 2013). This meant solving for an estimate of *W* by the following:

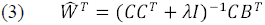

where λ is the ridge parameter penalty term, and *I* is a *k × k* identity matrix. We estimated an optimal λ for each voxel by evaluating Equation 3 over a range of λ values (0 to 750) for a balanced number of runs of the attention task (e.g., an equal number of runs from each attention condition). We then computed the Bayesian Information Criterion (BIC) for each of these λ values, estimating the degrees of freedom in the ridge regression as:

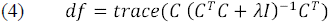

The λ with the smallest BIC was selected for each voxel. Since the attention comparisons are done within voxels, the varying λ penalty across voxels could not explain the attention effects we observed.

To select reliable voxels for analysis, we next implemented a set of conservative thresholding steps (Table 1). We first needed to select voxels with reliable visual responses, so we only kept voxels with trial beta weights that predicted at least 50% of the BOLD time courses in each scan session. Second, we only used voxels that could be successfully regularized with ridge regression. Any voxels with the maximum λ (750) were discarded, as this indicated that the ridge regression solution had not converged. Finally, we verified that the resulting regression model could predict an independent dataset, so we performed exhaustive leave-one-run-out cross validation for each attention condition. This ensured that the λ estimated across attention conditions produced reliable data for each condition separately. We estimated *W* using data from all but one run (Equation 3) and used this to predict the BOLD GLM trial estimate of the left-out run (Equation 2), separately for each condition. We then computed the mean correlation between the predicted & real BOLD GLM trial estimates across cross-validation folds for each voxel. Note that it is not possible to calculate a coefficient of determination on regularized data, since the process of ridge regression changes the scale of the predicted data (see supplemental discussion in Huth et al., 2012). We only kept voxels where this cross-validation r > 0.25 for all 3 conditions.

**Table 1.** VRF selection statistics, pooled across participants (N = 7)

| Region of interest | Total no. of localized voxels | No. of voxels after GLM thresholding | No. of voxels after regularizability threshold | No. of voxels after cross-validation threshold | No. of voxels after removing difference score outliers | Percent that survive all thresholds | RMSE fit error for surviving voxels |
| --- | --- | --- | --- | --- | --- | --- | --- |
| V1 | 3,723 | 3,540 | 2,438 | 989 | 931 | 25.01% | 0.1105 |
| V2 | 4,154 | 3,970 | 3,115 | 1,405 | 1,339 | 32.23% | 0.1087 |
| V3 | 3,698 | 3,519 | 2,839 | 1,520 | 1,435 | 38.81% | 0.0994 |
| V4 | 1,702 | 1,492 | 1,118 | 361 | 336 | 19.74% | 0.0783 |
| V3A/B | 1,988 | 1,922 | 1,440 | 443 | 416 | 20.93% | 0.0893 |
| IPS0 | 1,567 | 1,492 | 800 | 114 | 110 | 7.02% | 0.0882 |
| V1 – V4 | 13,277 | 12,521 | 9,510 | 4,275 | 4,041 | 30.44% | 0.1032 |
| V3A/B & IPS0 | 3,555 | 3,414 | 2,240 | 557 | 526 | 14.80% | 0.0894 |
| TOTAL | 16,832 | 15,935 | 11,750 | 4,832 | 4,567 | 27.13% | 0.1016 |

To quantify each vRF, we fit the spatial RF profile of each voxel with a smooth 2D function with 4 parameters: center, size, baseline, and amplitude (Fig 1b; Equation 2). Here, we define the vRF baseline as the voxel’s response that does not reliably depend on the position of the mapping stimulus (i.e., its constant offset). The vRF amplitude is defined as the spatially-selective increase in a voxel’s response above this baseline. Together, these two parameters index how much of the voxel’s response is due to a change in mapping stimulus position. Finally, the size and location parameters estimate the spatial selectivity and the spatial position preference of the vRFs, respectively. We first downsampled the vRFs by multiplying the estimated weights 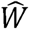 for each voxel (a 1 *× k* channel vector) by a smaller version of the spatial grid that contained the basis set (68 by 51 pixel grid; 10° by 5°). This speeded up the process of fitting the pixelwise surface with Eq. 2. This fitting process began with a coarse grid search that first found the best fit in a discrete grid of possible vRF parameters (center sampled in 1° steps over the mapped portion of the visual field; size constant logarithmically sampled at 20 points between FWHM of 10^0.01° and 10^1°). At each grid point, we estimated the best fit amplitude and baseline using linear regression. The grid point fit with the smallest root mean square error (RMSE) provided the initialization seed to a continuous error function optimization algorithm (fmincon in MATLAB). This fit had several constraints: the final solution must place the center within 2 grid points of the seeded fit (parameterized by position and size) and within the mapped visual field; the amplitude must be between 0 and 5; the baseline must be between -5 and 5 BOLD z-score units. Occasionally, this nonlinear fitting algorithm did not converge and resulted in a larger error than the original seed. In this case we took the best fit grid point as the final fit.

To test whether vRF fit parameters changed when participants focused spatial attention at different positions, we compared fits during each attend periphery condition with fits during the attend fixation condition. We computed a difference score (attend peripheral – attend fixation) to describe the magnitude of the attentional modulation. For example, a difference score of –2° in the FWHM of the vRF would indicate that the response profile width decreased when the participant attended to the peripheral target. This analysis revealed a subset of voxels with very large difference scores, which we determined to be due to noisy data or poor fits via manual inspection. Accordingly, we performed a final threshholding step for all vRF-based analyses: an elimination of outlier voxels with difference scores greater than three times the standard deviation of the population mean, where the population consists of the parameter difference scores for a given ROI (Table 1). After removing these outliers, we tested whether the vRF parameter difference scores differed significantly from 0 within a visual region of interest (ROI) by bootstrapping the distribution of difference scores across participants 10,000 times.

To determine if these vRF changes were modulated by their position in the visual field, we first calculated each vRF’s distance from the attended location (*v_dist_attn*) using its position during the fixation task. We then fit an *n*^th^ order polynomial to the vRF difference scores as a function of *v_dist_attn*, where *n* = 0, 1, or 2. This corresponds to a constant offset (0^th^ order), a linear fit (1^st^ order), or a quadratic or parabolic fit (2^nd^ order). These fits were cross-validated by fitting on 50% of the vRF difference scores and calculating goodness-of-fit (residual sum of squares and R^2^) on each of the 10,000 cross-validation iterations. These cross-validation iterations also provided confidence intervals on the coefficients for each polynomial. The most parsimonious fit was chosen by performing a nested F-test on the average residual sum of squares for each polynomial model.

We also tested whether vRF attentional modulations depended on hemisphere or visual hemifield, akin to the results reported for IPS0 – IPS5 in Sheremata and Silver, 2015. We sorted the voxels in each attention condition as contralateral or ipsilateral to the attended target. We then performed a series of non-parametric bootstrapped tests similar to a two-way ANOVA with attended hemifield and voxel hemisphere as factors. The vRFs were resampled with replacement across subjects 10,000 times. We then evaluated the two main effects and the interaction by computing a difference in the means of the groups or a difference in the slope between the group means, respectively. None of the tests for the effect of hemisphere and the interaction survived FDR correction, so we do not report those results here. We speculate that this null result is likely due to a lack of reliable voxels in anterior parietal cortex areas IPS1-5 in our study, where previous reports have found larger laterality effects (Sheremata and Silver, 2015).

### Population analysis (1): Fine spatial discriminability metric

To compute the spatial discriminability of a population of vRFs, we estimated the spatial derivative of each vRF at every point in the mapped visual field in 0.1° steps (Fig 1C). This was done by taking the slope of the vRF along the x and y direction at each pixel in the image of the visual field and squaring this value (Scolari and Serences, 2009, 2010). This measurement is a descriptor of how well a population code can discriminate small changes in the spatial arrangement of the stimulus array, which depends on the rising and falling edges of a tuning curve rather than the difference between the peak response and a baseline response (Regan and Beverley, 1985; Pouget et al., 2003; Butts and Goldman, 2006; Navalpakkam and Itti, 2007; Scolari and Serences, 2009, 2010). To restrict our measurements to the relevant area near the peripheral target, we computed discriminability values within 1 degree of the center of each target across both spatial dimensions (x and y). These were summed and divided by the maximum discriminability value in that population to make the results comparable despite changes in vRF coverage or responsiveness.

### Population measurements (2): Stimulus reconstructions using an inverted spatial encoding model

In addition to computing the discriminability metric described above, we also reconstructed an image of the entire visual field on each trial using a population-level encoding model. Compared to the local spatial discriminability index, this is a more sensitive method of assessing the amount of spatial information encoded in an entire population of voxels because it exploits the pattern of response differences across voxels, rather than treating each voxel as an independent encoding unit (Serences and Saproo, 2012; Sprague et al., 2015).

We trained the spatial encoding model using a procedure similar to the vRF estimation analysis described above (Fig 4a). This yields an estimated matrix of weights, 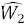, which specifies how much each voxel in a region of interest responds to each of the spatial channels (Brouwer and Heeger, 2009; Serences and Saproo, 2012; Sprague and Serences, 2013; Sprague et al., 2015). We then solved Eq. 1 using the Moore-Penrose pseudoinverse with no regularization:

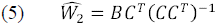

*C* was constructed using a set of 54 evenly tiled spatial filters (Eq. 2; 9 horizontal / 6 vertical; spaced 1.25° apart; 1.56° FWHM). 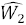 was estimated using the data from the jittered position runs. This was done separately for each participant, using a training set balanced across the conditions of interest (e.g., an equal number of attend left and attend right runs and all attend fixation runs, since fixation is the neutral condition).

**Figure 2.**
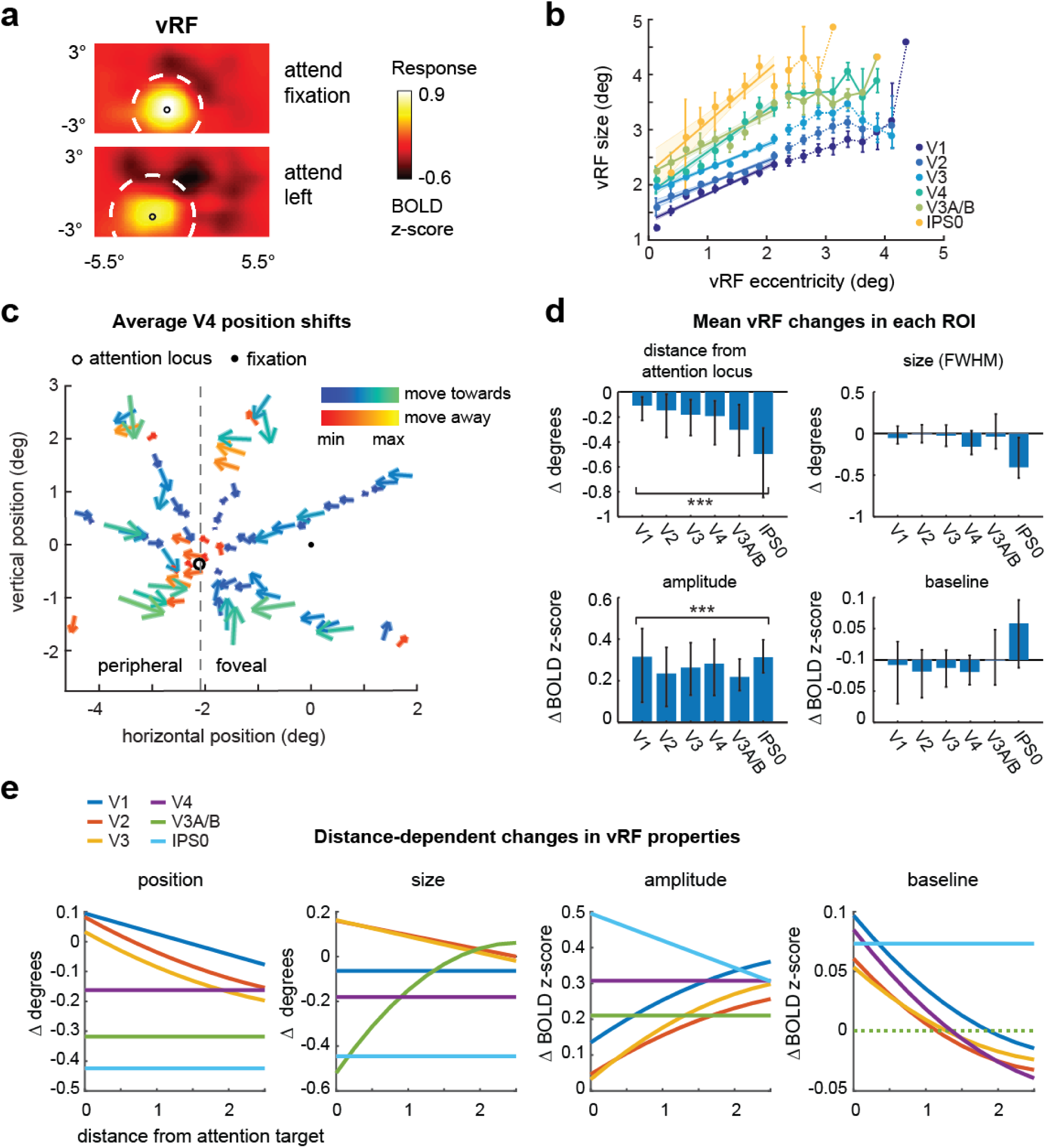
Changes in voxel receptive fields (vRFs) across attention conditions. We separatelyestimated vRFs for every voxel in visual and posterior parietal areas, discarding poorly estimated or noisy voxels (Table 1). Unless otherwise specified, figure data is averaged across subjects and error bars show 95% confidence intervals computed by resampling the data distribution. (**a**) An example vRF shows that attending covertly to the left location shifts the center of the receptive field profile to the left, when compared to the neutral attend fixation condition. Voxel is fromsubject AR in area V3A/B. (**b**) Our vRF estimates reproduced the canonical size-eccentricity relationship (positive slope in all ROIs, p < minimum possible p-value, e.g., 1/10000 iterations) and the increase in slope between visual regions. (**c**) Preferred position changes of V4 vRFs with covert spatial attention. We binned each vRF by its position during the attend fixation condition. The origin of each arrow is the center of each position bin. The end of the arrow shows the average position shift of the vRFs within that position bin during the attend peripheral conditions (left/right are collapsed and shown as attend left). The majority of vRFs shift toward the attended location (blue-green color map vs. red-yellow). (**d**) Mean changes in vRF parameters (attend peripheral target – attend fixation) in each visual area. (**e**) Attentional modulations of each vRF parameter plotted by the vRF’s distance from the attention target computed from its position during the attend fixation task (Table 2).

**Figure 3.**
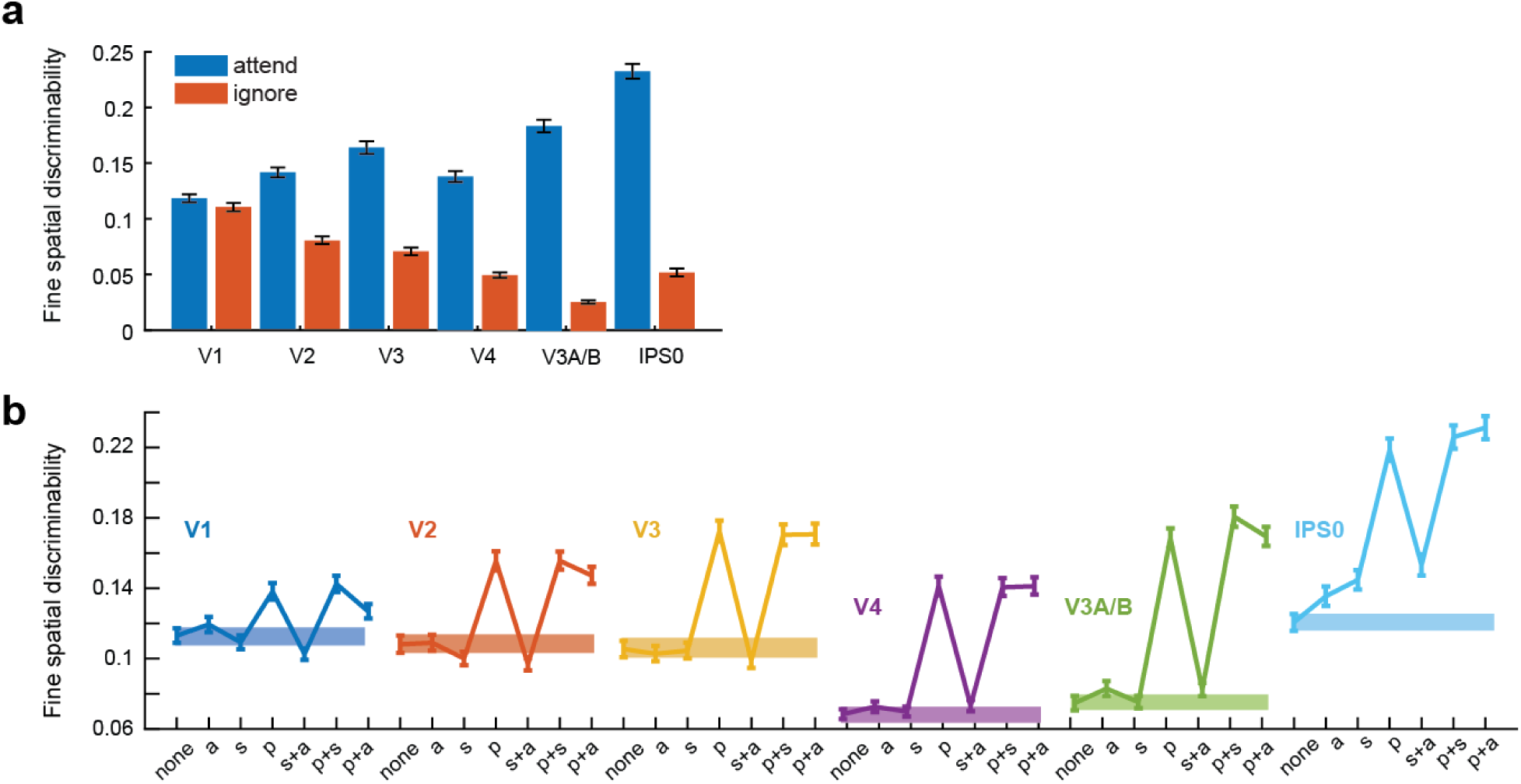
Spatial discriminability increases with attention and is mediated by position changes in vRFs. Error bars depict bootstrapped 95% CIs. (**a**) We formulated a measurement to describe the ability of a population of voxels to make fine spatial discriminations around the attention target. We used the properties of each voxel’s spatial tuning curve to make this measurement (**Materials and Methods**). Spatial discriminability increased when subjects attended the target, compared to when they ignored the target in the opposite hemifield (resampled p < minimum possible p-value (1/1000) for all ROIs for all ROIs). (**b**) The discriminability metric was recomputed for vRFs with a variety of attentional modulations. (*none* = vRF parameters during the neural attend fixation condition; *a* = amplitude; *s* = size; *p* = position). Spatial discriminability increased significantly when we applied position changes measured during the attend L/R task to the vRFs compared to when we applied no parameter changes (solid bar). By contrast, applying size changes did not change spatial discriminability in most ROIs, although it did cause a small increase in IPS0.

**Figure 4.**
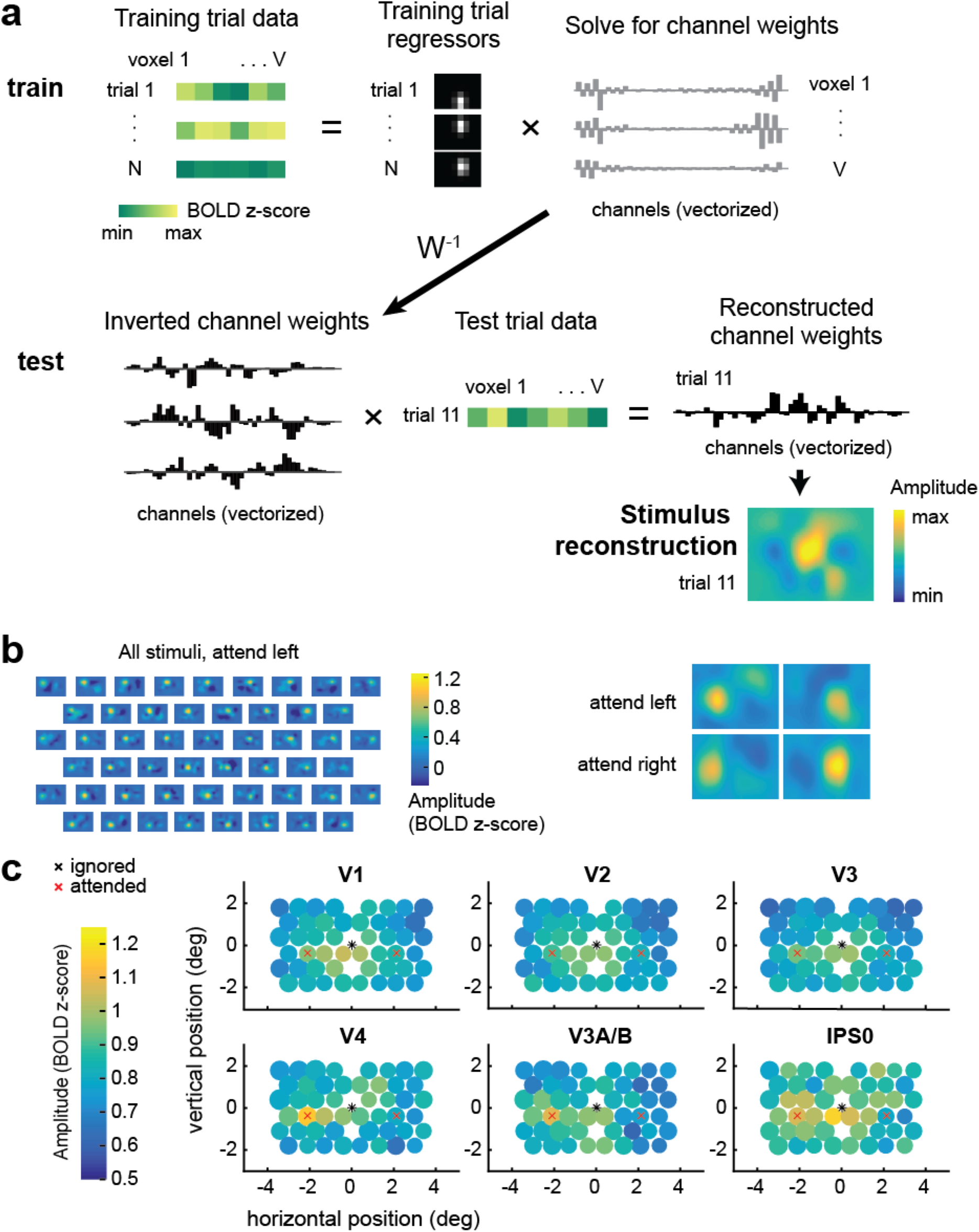
Multivariate inverted encoding model (IEM) used to reconstruct the mapping probe stimuli. (**a**) To train the IEM, we first take the BOLD data from all voxels within a visual region from a subset of training trials. Then, we solve for a set of channel weights using least squares regression. To reconstruct the stimulus, we invert this weight matrix and multiply it with BOLD data from the same voxels during a test trial. This yields a reconstructed channel response profile, which can be transformed into a reconstruction of the mapping stimulus on every trial in each attention condition. Data shown are examples from participant AR for a subset of V1 voxels. (**b**) Example stimulus reconstructions for participant AI, V1. These reconstructions were averaged across trials with the same position, yielding 51 reconstructions – one for each unique position in the test dataset. In the left panel, the same averaged position reconstructions are shown for each condition. The amplitude on the left is higher when attending left, and on the right when attending right. (**c**) Average reconstruction sizes and amplitudes for each stimulus position (collapsed across condition; left is attended). The diameter of the circle depicts the average fit FWHM of the reconstructions at that spatial position. Reconstruction amplitude was greater in the attended hemifield compared to the ignored hemifield in areas V3A/B and V4 (p <= 0.005; Table 3; Figure 5).

**Figure 5.**
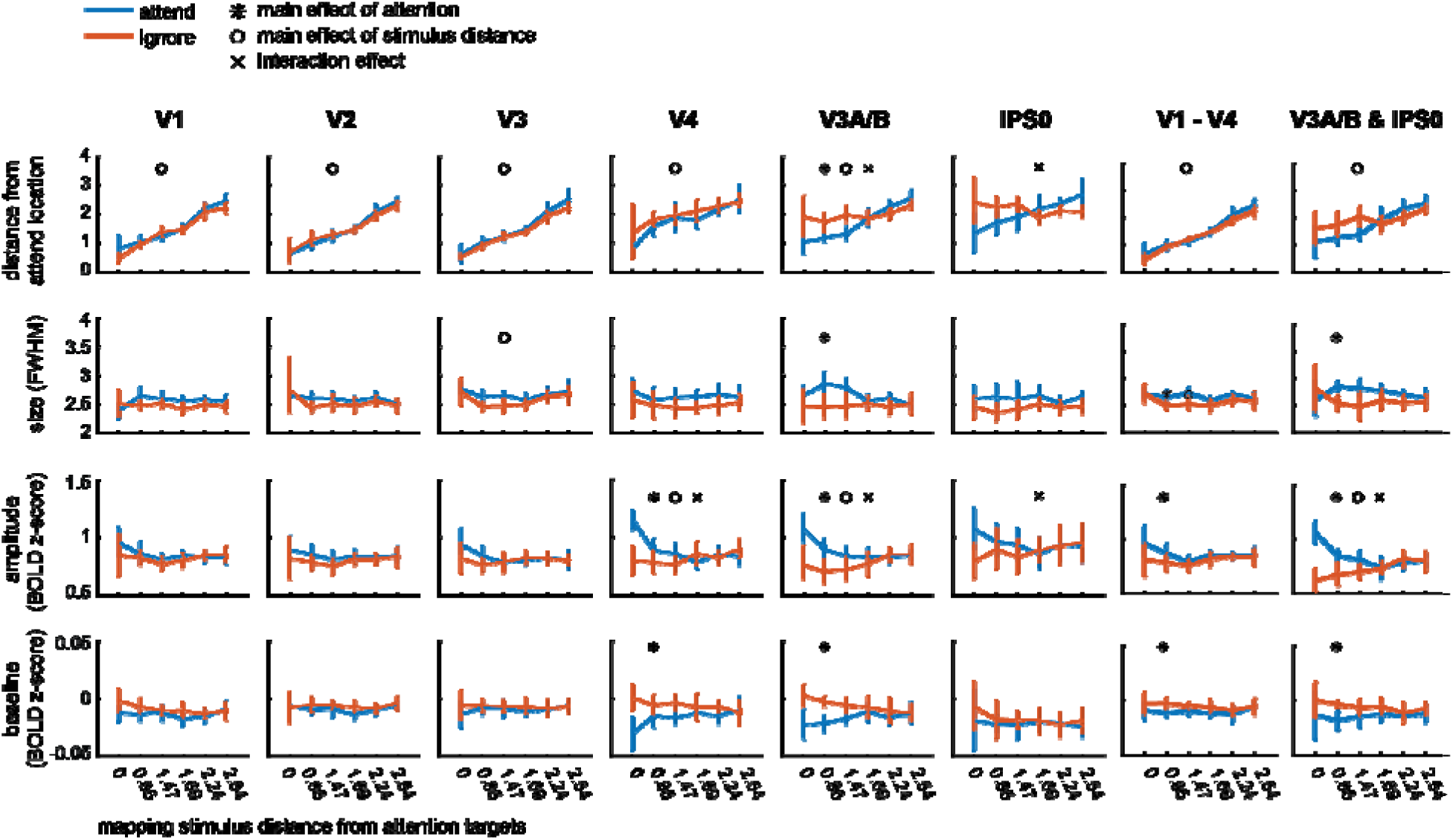
Reconstruction parameters as a function of mapping stimulus distance from the covertly attended locations and attention hemifield (attended vs. ignored). See Table 3 for complete list of p-values.

**Table 2.** Mean coefficients for polynomial fits of how vRF parameter change is modulated bydistance from the attended location (*v_dist_attn*)

|  | Position | Size | Amplitude | Baseline |
| --- | --- | --- | --- | --- |
| V1 | -.069, .095 | -.064 | -.019, .137, .135 | .011, -.073, .010 |
| V2 | .015, -.133, .082 | -.064, .160 | -.016, .125, .046 | .011, -.066, .061 |
| V3 | .017, -.135, .033 | -.073, .163 | -.025, .170, .032 | .009, -.053, .054 |
| V4 | -.162 | -.181 | .308 | .011, -.078, .085 |
| V3A/B | -.318 | -.091, .461, -.520 | .210 | <.001 |
| IPS0 | -.425 | -.445 | -.076, .495 | .073 |

<sup>a</sup> Number of reported coefficients in the table correspond to the polynomial order which was yielded the most parsimonious fit to the data (e.g., 1 coefficient for $n = 0$ , 2 coefficients for $n = 1$ , etc.).

**Table 3.** 2-way ANOVA results for reconstruction parameter changes (*s_dist_attn*x attentionhemifield).

|  | V1 | V2 | V3 | V4 | V3A/B | IPS0 | V1 – V4 | V3A/B & IPS0 |
| --- | --- | --- | --- | --- | --- | --- | --- | --- |
| Omnibus test |  |  |  |  |  |  |  |  |
| Position | **<.001** | **<.001** | **<.001** | **<.001** | **<.001** | **.001** | **<.001** | **<.001** |
| Size | .216 | .565 | .019 | .428 | **.006** | .121 | **.001** | .110 |
| Amplitude | .174 | .579 | **.024** | **<.001** | **<.001** | **.008** | **.016** | **<.001** |
| Baseline | .088 | .734 | .934 | **<.001** | **.001** | .937 | **.015** | .241 |
| Main effect of distance |  |  |  |  |  |  |  |  |
| Position | **<.001** | **<.001** | **<.001** | **<.001** | **<.001** | .192 | **<.001** | **<.001** |
| Size |  |  |  |  | .484 |  | **.019** |  |
| Amplitude |  |  | .140 | **.002** | **.005** | .478 | .100 | **.002** |
| Baseline |  |  |  | .829 | .916 |  | .210 |  |
| Main effect of attention |  |  |  |  |  |  |  |  |
| Position | .371 | .916 | .346 | .067 | **.005** | .254 | .401 | .343 |
| Size |  |  |  |  | **.003** |  | **.005** |  |
| Amplitude |  |  | .069 | **<.001** | **.005** | .158 | **.049** | .004 |
| Baseline |  |  |  | **.001** | **<.001** |  | **.004** |  |
| Interaction of distance & attention |  |  |  |  |  |  |  |  |
| Position | .052 | .588 | .541 | .657 | **<.001** | **<.001** | .121 | .026 |
| Size |  |  |  |  | .077 |  | .271 |  |
| Amplitude |  |  | .064 | **<.001** | **<.001** | **.004** | .224 | **<.001** |
| Baseline |  |  |  | .019 | .011 |  | .370 |  |

<sup>a</sup> **bold** numbers indicate that the p-value passed FDR-correction ( $q = .05$ , corrected across ROIs and comparisons within each parameter).

**Table 4.** RMSE (and 95% CIs) between reconstructions from the reduced dataset (only using voxels with RFs) or from different versions of the layered IEM using the same voxels.

|  | **Reduced data** | **p/s/a/b** | **p/a/b** | **p/s/b** | **s/a/b** | **p/a** | **s/a** | **p/s** | **p** | **a** | **s** | **none** |
| --- | --- | --- | --- | --- | --- | --- | --- | --- | --- | --- | --- | --- |
| Combined occipital V1 – V4 | 0.133<br>[0.109, 0.170] | 0.146<br>[0.146, 0.146] | 0.148<br>[0.147, 0.148] | 0.142<br>[0.141, 0.143] | 0.194<br>[0.194, 0.194] | 0.148<br>[0.148, 0.148] | 0.194<br>[0.194, 0.194] | 0.141<br>[0.141, 0.141] | 0.143<br>[0.143, 0.143] | 0.194<br>[0.194, 0.194] | 0.194<br>[0.194, 0.194] | 0.193<br>[0.193, 0.193] |
| Combined parietal V3A/B & IPS0 | 0.834<br>[0.744, 0.959] | 0.415<br>[0.410, 0.419] | 0.426<br>[0.422, 0.431] | 0.411<br>[0.407, 0.416] | 0.443<br>[0.439, 0.447] | 0.425<br>[0.421, 0.430] | 0.451<br>[0.447, 0.456] | 0.410<br>[0.405, 0.416] | 0.410<br>[0.406, 0.414] | 0.441<br>[0.437, 0.445] | 0.447<br>[0.442, 0.452] | 0.834<br>[0.744, 0.959] |

<sup>a</sup> To generate CIs, the resampling of the real data is performed at the level of the fits to the reconstructions, whereas resampling layered IEM RMSEs is described in **Materials and Methods**

To reconstruct a representation of the mapped visual space, we inverted the model and multiplied the pseudoinverse of the estimated weight matrix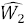 with a test dataset from the discrete position runs (*B*_2_), yielding estimated channel activations for each trial (*C*_*2*_; *k*_*2*_ channels by *t* test trials) (Equation 6). Thus, we refer to this analysis as the inverted encoding model (IEM).

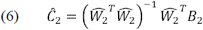

Because of mathematical constraints on inverting *W*_2_ (number of voxels must be greater than number of channels), we included all voxels in each ROI instead of just the subset of well-fit voxels used in the vRF analyses described above. We computed Eq. 6 twice using different test datasets, once for the discrete position attend left runs and once for the discrete position attend right runs.

When we multiply the resulting channel activations by a grid of pixels that define the spatial channels, we obtain a spatial representation of the entire visual field on each trial. This image contains a stimulus reconstruction showing where the checkerboard should have been given the trained model and the activation pattern across all voxels in the independent test set. The stimulus reconstructions were then fit in the same manner as the vRFs, using Eq. 1 to estimate the center, size, amplitude, and baseline (mean fit RMSE across all ROI reconstructions 0.114; 95% CI [0.102, 0.312]). Here, the baseline is an estimate of the multivariate reconstruction that is spatially non-selective—i.e., not significantly modulated by the position of the mapping stimulus. The amplitude describes the maximal increase in that reconstruction relative to baseline when the mapping stimulus is on the screen.

To assess how attention changed reconstructions of the mapping stimulus across the visual field, we first computed a difference score that described the effect of attention by folding the visual field in half (i.e. collapsing across hemifield) and comparing parameters in the attended vs. ignored hemifield. We excluded the reconstructions that fell along the vertical meridian (3 of 51 stimulus positions). This allowed us to control for the overall effect of eccentricity while remaining sensitive to other spatial patterns in stimulus reconstruction modulations.

We then set up a single factor repeated measures omnibus ANOVA to determine which pairs of ROI and parameter (e.g., V1 size, V1 amplitude, etc.) were affected by either attention or Euclidean distance from the target stimuli. The attention factor had two levels (attend/ignore) and the distance factor had 6 levels (6 evenly spaced distance bins from 0° to 2.54°). Based on the results of this omnibus test, we tested any significant ROI-parameter combination in a 2-way repeated measures ANOVA of attention by distance. To estimate the p-values for these tests, we generated empirical null distributions of the F-scores by randomizing the labels within each factor 10,000 times within each participant. Reported p-values are the percentage of the randomized F-scores that are greater than or equal to the real F-scores.

### Population analysis (3): Layered spatial encoding model to link vRFs to multivariate stimulus reconstructions

In order to test how changes in the response properties of the underlying vRFs contributed to changes in the fidelity of region-level stimulus reconstructions, we generated simulated patterns of voxel activity on every trial by predicting the response to each stimulus based on the vRF fit parameters. We then used this simulated data to estimate and invert a population-level spatial encoding model, as described above (Fig 6a).

**Figure 6.**
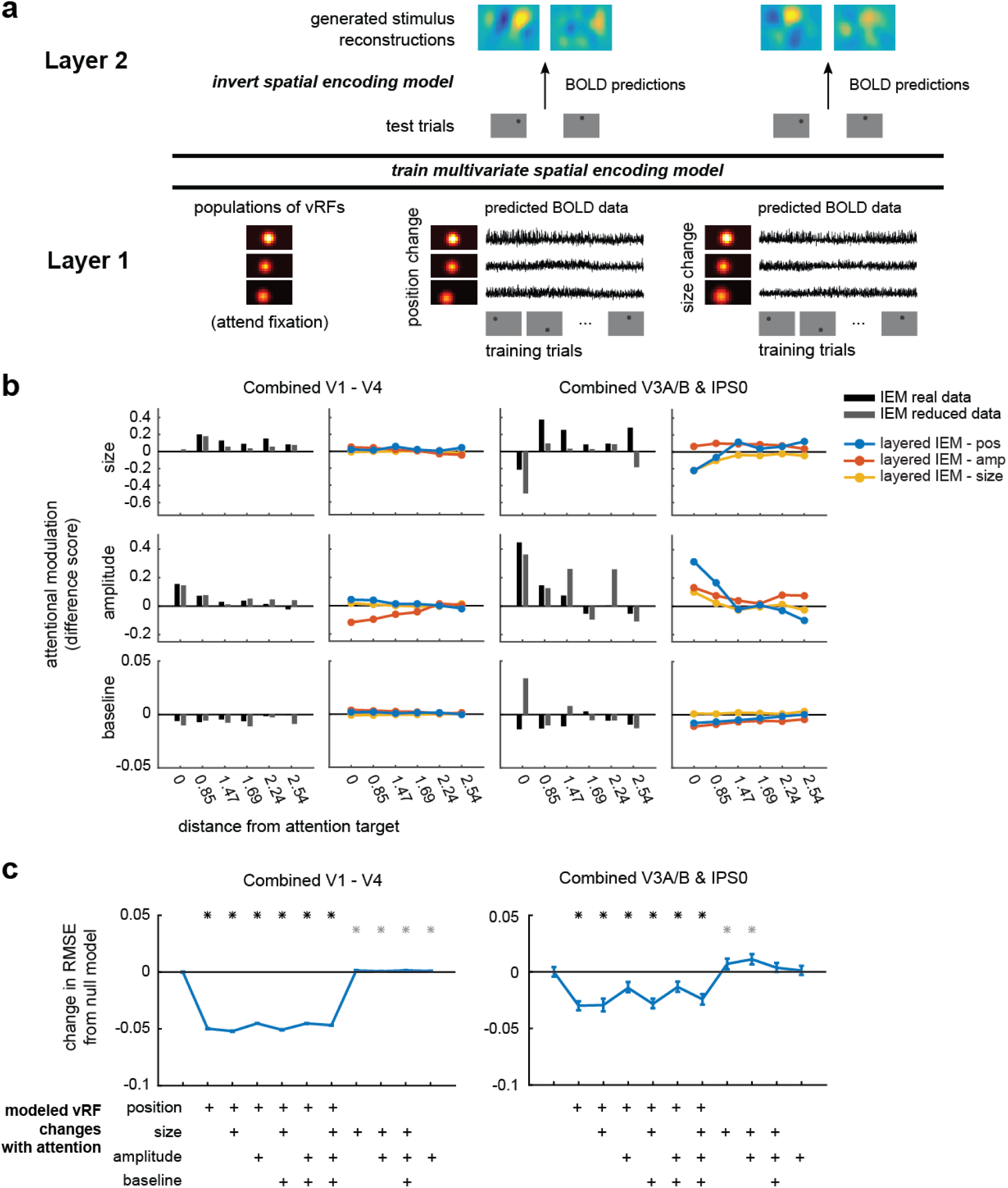
A layered spatial encoding model reveals how different sets of vRF changes lead to enhancements in multivariate stimulus reconstructions. (**a**) The first layer of the model uses the vRF fits to generate BOLD data from every subject’s real trial sequence. Then the BOLD data from all voxels within one ROI is used to train a multivariate spatial encoding model and reconstruct the mapping stimuli. (**b**) Change in reconstruction amplitude in the attended vs. theignored hemifield. We only show reconstruction parameters with significant attentional modulations in the prior IEM analysis (Fig 5, Table 3). Stimulus reconstructions computed with a reduced number of voxels (gray bar) largely reproduce the pattern of attentional modulations observed in IEMs computed with all voxels (black bar). Furthermore, a comparison of layered IEMs using simulated data revealed that vRF position changes (blue lines) in the first layer of the model are better at reproducing the amplitude modulations in the stimulus reconstructions of the parietal ROI than models which simulate changes in vRF size or amplitude (yellow & red lines). (**c**) RMSE between each set of IEM fits and the full empirical dataset fits shown in Fig 5. The null baseline model (far left) is a layered IEM where the vRF parameters are the same across all attention conditions. We then added vRF attentional modulations for each parameter as shown in the matrix below the plot, where all models with position changes are on the left side. # indicate an FDR-corrected p-value <.05 for models that differed significantly from the null baseline model. Gray asterisks indicate an increase in RMSE from the null model, whereas black asterisks indicate a decrease.

Note that for these simulations, we could only use well-fit voxels to generate simulated BOLD timeseries. This constrained the analysis to ROIs with at least as many vRFs as spatial filters used to estimate the spatial encoding model. To ensure that we could include most participants in the layered encoding model analysis, we created two large ROIs by merging the smaller retinotopically defined regions described above. The occipital ROI consisted of V1, V2, V3, and V4 defined for each subject. The posterior parietal ROI consisted of V3A/B and IPS0. The vRFs in the parietal ROI show distinct patterns of attentional modulations (Fig 2e), suggesting that V3A/B and IPS0 are both anatomically and functionally distinct from the occipital regions (see also de Haas et al, 2014). Although merging ROIs increased the number of voxels available for the encoding model analysis, we still did not have enough voxels in the parietal ROI to estimate the layered encoding model for 3 of the 7 participants (AL, AR, AU). However, the remaining data from 4 participants were sufficient to produce stable, subject-averaged results.

To simulate each voxel’s BOLD response on every trial that the participant completed in the real experiment, we first created a high-resolution set of spatial channels (21 by 11 channels spaced 0.5° apart, FWHM = 0.65°) and generated weights for each channel based on the vRF fit obtained from prior analysis. That is, we evaluated Eq. 2 for each channel at the vRF’s fit center and adjusted the response gain by multiplying this result by the fit amplitude and adding the fit baseline. We then added independent Gaussian noise to each of these channel weights, simulating a small amount of variance in the voxel’s response (σ = 0.5). Each voxel’s channel weights were then multiplied by the idealized channel response on each trial (that is, the channel filter convolved with the stimulus mask), effectively simulating the BOLD response on each trial for the entire population of voxels based on their measured vRFs. We added Gaussian noise to this simulated response as well (σ = 0.5). We then computed stimulus reconstructions using the same method as described above (the IEM in **Population analysis (2)**), averaging resulting reconstructions across participants and like positions before fitting.

To ensure the stability of the reconstructions that were based on simulated data, we repeated the simulations 100 times and averaged across the fits of all iterations to generate the plots in Fig 6b. Then, to compare how well the layered model reproduced the attentional modulations observed in stimulus reconstructions generated with real data, we calculated an error metric between the layered IEM and the real data. We first calculated reconstruction difference scores across attention condition (attended – ignored; see **Population analysis (2)**). This yielded 24 difference scores each for both attention conditions in both the layered IEM data and the empirical data. Since the empirical data did not have any repeated iterations, we averaged across all 100 iterations of the layered model to match the dimensionality of the real reconstructions (2 conditions x 24 difference scores x 4 parameters). We could then calculate the root mean square error (RMSE) between the difference scores from the full empirical dataset (i.e. the data shown in Fig 5) and the modeled data. This was used as a metric to describe the goodness-of-fit of each layered IEM.

We then tested how different vRF attentional modulations contributed to changes in the population-level stimulus reconstructions. To test how shifts in vRF centers contributed to population-level information, we modeled voxels that had the same fit center across both attention conditions, simulated their BOLD responses on each trial, and generated stimulus reconstructions from these data. The voxel’s vRF center was defined as the vRF center fit from the neutral attend fixation data. A similar procedure was repeated for all reported combinations of parameter changes across conditions. Again, whichever parameter was held constant took its value from the neutral attend fixation condition.

To calculate the confidence intervals on the RMSE changes in Fig 6c, we resampled with replacement across the 100 model iterations and refit the average across these 100 instances. This resampling procedure was repeated 500 times to generate a distribution of fits to the model data. We then took the difference between the RMSE of the null model, in which no parameters varied between attention conditions, and the RMSE of the model which held some number of vRF parameters constant across attention conditions.

## RESULTS

### Modulations of vRF properties with spatial attention

We estimated single voxel receptive fields (vRFs) for each voxel in 6 retinotopically-identified visual areas from V1 to IPS0. The estimation of vRFs was done independently for each attention condition so that we could compare a single voxel’s spatial tuning across conditions.

To confirm that the fit sizes were consistent with previous results, we fit a line to the estimated sizes as a function of the vRF center eccentricity. First, we combined all vRFs across participants and conditions in each ROI. We then binned the vRF centers every 0.25° from fixation and calculated the mean size (Fig 2b). We first replicated an increase in vRF size with increasing eccentricity, and an increase in the slope of this relationship across visual regions (Gattass et al., 2005; Dumoulin and Wandell, 2008; Amano et al., 2009; Harvey and Dumoulin, 2011) (Fig 2b). These observations confirm that our method produced reasonable vRF estimates that were consistent with previous reports.

Next, we examined how covert attention to the peripheral attention targets modulated vRF properties, relative to the attend fixation condition. Overall, the center position of vRFs shifted significantly closer to the attended location (p < 0.005 in all ROIs, Fig 2d). This finding is consistent with previous reports in humans and in monkeys for both covert attention tasks and saccade tasks (Womelsdorf et al., 2006, 2008; Klein et al., 2014; Zirnsak et al., 2014).

While we did observe changes in the size of individual vRFs, the mean change was not significantly different from zero (p > 0.05 in all ROIs). Size increases have been previously reported in tasks that required subjects to attend to the mapping stimulus, which moved on each trial (Sprague and Serences, 2013; Kay et al., 2015; Sheremata and Silver, 2015). Accordingly, if attention causes the center of RFs to shift toward the attended target, these combined shifts in position would average out to form a larger RF estimate. In contrast, mapping vRFs while maintaining a fixed locus of attention would nullify the size increase, consistent with the results we observed (Fig 2d). Another study which also found increases in vRF size with attention required subjects to attend the fixation point while they manipulated the perceptual load, or difficulty, of the attention task (de Haas et al., 2014). In our study, we intentionally kept task performance constant and could not evaluate effects of difficulty on the parameters of vRFs.

We also found an overall increase in vRF amplitude with attention (p < 0.001 for all tests). Since these measures were calculated relative to a fixation task, these data suggest that covert spatial attention to a peripheral location caused widespread position and gain modulations in all vRFs across the visual field.

It is unclear whether these attentional modulations are limited to areas near the attended target, or whether they are uniform across the visual field. For example, vRF position shifts could result in a radial convergence of RFs towards the attended target, or a uniform shift of RFs along a vector extending from fixation to the attention or saccade target (Tolias et al., 2001; Klein et al., 2014; Zirnsak et al., 2014). Furthermore, reports of other RF properties (such as size) modulating with attention have been mixed (Connor et al., 1996, 1997; Womelsdorf et al., 2008; Niebergall et al., 2011; Sprague and Serences, 2013; de Haas et al., 2014; Klein et al., 2014; Kay et al., 2015; Sheremata and Silver, 2015). We therefore examined whether each of the vRF parameter changes was dependent on the vRF’s location in the visual field, relative to the attended location. First, we created radial distance bins centered on the left or right attended locations, and sorted voxels into these bins based on their preferred position during the fixation condition. After this sorting procedure, data from the right condition were flipped and collapsed with the left condition.

When we plotted vRF position changes in each bin, we found that spatial attention caused vRF position shifts that converged on the attended location (two-tailed sign test on vector direction, p < .001 in all ROIs). That is, vRFs shifted closer to the attended location (Fig 2c), compared to when subjects attended fixation (mean shift across all vRFs and ROIs: -0.239°, 95% C.I. [-0.566, -0.048], Fig 2d). Note that small eye movements toward the attended location cannot explain receptive field convergence: this would cause all vRFs to shift in the same horizontal direction, rather than radially converging on one point. These data are consistent with results from both humans (Klein et al., 2014) and macaques (Connor et al., 1996, 1997, Womelsdorf et al., 2006, 2008) that use a similar task. However, the prior study in humans focused only on vRFs with preferred locations that were foveal to the attended location, and the studies in macaques only report RF position changes in V4 and MT. By contrast, our data show that vRF centers converge on the attended location across all visual areas, including primary visual cortex, and that this pattern of modulations includes vRFs peripheral to the attended target.

These plots (Fig 2a, 2d) also suggested that vRFs farther from the attended location underwent larger position changes than vRFs near the attended location. That is, the magnitude of the attentional modulation may be dependent on the distance between the vRF and the attended target. To test for this, we fit a polynomial to the vRF parameter changes as a function of distance from the attended location (**Materials and Methods**). We selected the most parsimonious fit ranging from a mean change in vRF parameter (0^th^ order polynomial) to a parabolic change (2^nd^ order polynomial) by conducting a nested F-test (Table 2). The best polynomial fits are plotted in Fig 2e.

This analysis allowed us to characterize trends in vRF attentional modulations across space. Note that it also implicitly tests whether voxels contralateral to the attended target respond differently than ipsilateral voxels. This is because vRFs near the attended target will mostly originate from the contralateral side of visual cortex. Therefore, any fit lines with a significant slope imply there is a difference between contralateral and ipsilateral voxels (Sheremata and Silver, 2015). A separate test described in **Materials and Methods** confirmed that contralateral voxels differed significantly from ipsilateral voxels in the areas where we saw the highest fit slopes in Fig 2e (FDR-corrected p <.05 for position: V1, V2; size: V3; amplitude: V3, IPS0). However, since the fit lines illustrate how these changes occur over space, we discuss those data here instead (Fig 2e).

In early visual areas V1 through V3, vRFs near the attention target were slightly repelled from the target, whereas vRFs farther from the target were attracted towards the target. In later visual areas, vRFs were uniformly attracted towards the attention target. We saw a similar pattern of results with size modulations: early visual areas showed an increase in vRF size near the attention target, and decreased size farther away. However, in areas V3A/B and IPS0, vRF size decreased near the attention target.

The pattern of vRF amplitude modulations was also segregated between the early and later visual areas. All vRFs increased in amplitude with attention, but the slope of this relationship inverted from early to later visual areas. In V1 – V3, the slope if positive, such that voxels ~2° away from the attention target increase in amplitude more than voxels right at the target position. The amplitude increase is constant in V4 and V3A/B. Finally, in IPS0, the slope inverts to become negative, so that voxels near the attention target increase in amplitude more than voxels farther away. Lastly, we found an increase in vRF baseline near the attended target in V1 – V4, but a uniform increase in baseline in IPS0. Overall, we found that the type and magnitude of the attentional modulation in different visual areas changes as a function of the spatial relationship between vRFs and the attended target. This is consistent with findings from macaque neurophysiology, which had suggested that amplitude and size changes depend on where the RF is located in relation to the attended target (Connor et al., 1996; Niebergall et al., 2011).

Note that these fits only describe the gain modulations with respect to the voxel’s position during the attend fixation task. However, these parameter changes likely interact with one another, such that a voxel that shifts toward the attended location will also increase in amplitude. Hence, to determine how the joint patterns of vRF modulations change the spatial information content of a representation, in the next section we discuss two different population-level measures that combine data across the population of vRFs in each ROI.

### Increases in spatial discriminability depend primarily on vRF position shifts

Next, we assessed how different types of RF modulations influenced the precision of population-level codes for spatial position. We first computed a discriminability metric that described the ability of a population of tuning curves (here, voxel receptive fields) to support fine spatial judgments (**Materials and Methods**). When we computed this metric based on the measured vRF properties from each condition, spatial discriminability near the attended target increased relative to the ignored target in the opposite visual hemifield (Fig 3a).

We then tested how different types of vRF modulations (such as size changes or position shifts) affected this spatial discriminability metric. As a baseline comparison, we first computed discriminability based on vRFs estimated during the attend fixation runs for each subject. We then added different sets of observed attentional modulations to the population before recomputing spatial discriminability. For example, we shifted all the vRF centers to match the measurements when a subject was attending to the left target and computed discriminability near the attended target. Since the response baseline of a vRF does not affect the discriminability metric, we excluded this type of attentional modulation from these analyses.

Across all ROIs, we found that vRF position shifts played the biggest role in increasing fine spatial discriminability compared to changes in size or changes in amplitude (Fig 3b). Position modulations alone led to a large increase in spatial discriminability, while other combinations of parameter modulations only had an impact if we added in position shifts (i.e. a change in size and position increased discriminability, but size alone did not). The only departure from these patterns was observed in IPS0, where all attentional modulation types increased spatial discriminability, but position changes increased spatial discriminability the most.

### Spatial attention increases the fidelity of population-level stimulus reconstructions

By design, the spatial discriminability metric we computed is only informative about local spatial representations, and cannot assess how different patterns of vRF modulations might result in representational changes across the visual field. To address this point, we built a multivariate spatial encoding model to measure how attention changes the representations of visual information in disparate parts of space. This also allowed us to further test the effects of vRF modulations on the encoding properties of the population, including response baseline changes that were not captured by our discriminability metric.

The spatial inverted encoding model (IEM) reconstructed an image of the entire visual field on each test trial. We first trained the model using the responses of each voxel on a set of training trials with known mapping stimulus positions. We then created image reconstructions on independent test trials by inverting the model and multiplying it by the voxel responses during each test trial (Fig 4a; **Materials and Methods**). Each image contained a representation of where the mapping stimulus should have been given the pattern of voxel activations on that particular trial. The IEM successfully reconstructed the task-irrelevant mapping stimuli using activation patterns across voxels in each visual area from V1 through IPS0 (Fig 4b; grand mean error between fit and actual position 2.40°, 95% CI [0.55°, 4.97]).

We used these stimulus reconstructions as a proxy for the quality of the spatial representations encoded in a population of voxels. This is line with previous studies showing that stimulus reconstructions change in amplitude or size as a function of cognitive demands. (Brouwer and Heeger, 2013; Ester et al., 2013; Sprague and Serences, 2013; Sprague et al., 2014, 2015, 2016).

First, we compared how reconstructed representations of each mapping stimulus changed as subjects shifted their spatial attention. We ran a repeated measures ANOVA of *attention* x *distance bin* for each reconstruction fit parameter (**Materials and Methods**). Here, a main effect of attention would suggest that stimulus reconstructions in the attended hemifield changed in a consistent way compared to the ignored hemifield. A main effect of distance would suggest that stimulus reconstruction changes had a consistent spatial pattern across both the attended and ignored hemifields. This would occur when a stimulus’ representation was altered with distance from the attention target. For example, the stimulus reconstruction center should vary linearly with the stimulus’ true distance from the attention target. And lastly, an interaction effect would suggest that the distance effect was dependent on whether the reconstruction belonged to the attended or ignored hemifield. In our task, the reconstructed stimuli are always irrelevant to the task of the observer. We therefore predicted an interaction effect where spatial attention would selectively modulate stimulus reconstructions within the hemifield of the attended location, but not the opposite hemifield (Connor et al., 1996, 1997).

We found that reconstruction amplitude was selectively increased near the attended location in V4, V3A/B, and IPS0 (interaction effect, bootstrapped p < 0.005; Fig 5; Table 3). This can be interpreted as a local boost in SNR. Prior reports found that attending to the mapping stimulus – as opposed to attending to a peripheral target in the current experiment – caused an increase in the amplitude of all stimulus reconstructions (Sprague and Serences, 2013). That is, representations of task-relevant stimuli increased in SNR. We find here that even representations of task-*irrelevant* stimuli near the attended region of space increase in amplitude, consistent with the idea of an attentional ‘spotlight’ which boosts the fidelity of spatial representations near the attention target.

Although the amplitude interaction effect was present in most visual areas we tested (Fig 5), we found other effects limited to V3A/B and IPS0 that involved modulations in stimulus representations in the ignored hemifield. In these regions, we found that stimulus reconstructions in the ignored hemifield shifted away from the ignored target location (interaction, bootstrapped p < 0.005). We also observed a relative size increase near the ignored attention stimulus in IPS0 (interaction, bootstrapped p < 0.005). These results suggest that stimulus reconstructions in the ignored hemifield are less spatially precise in posterior parietal cortex. Finally, there was also a main effect of attention on reconstruction size and baseline in areas V3, V4 & V3A/B (bootstrapped p’s <= 0.005). However, unlike the interaction effect in IPS0, these size and baseline changes did not vary as a function of distance between the reconstruction and the attended target location.

### Using a layered encoding model to explore how single voxel RFs change population-level codes

In our final analysis, we used a layered spatial encoding model to determine how changes in vRF properties affected the representations of mapping stimuli in the multivariate reconstructions discussed in the previous section (Fig 1c; Fig 4a). The goal of this analysis was to determine which vRF modulations contribute the most to the observed increase in the amplitude of stimulus representations around the attended location (Fig 5). This analysis thus complements our analysis of the spatial discriminability metric which demonstrated that vRF position changes significantly increased the ability of the population to make fine spatial discriminations near the attention target (Fig 3c).

The layered spatial encoding model we built links the response properties of single voxels to the encoding properties of a whole population of voxels in a region of visual cortex (Fig 6a). In the first layer of the model, we used the fit vRFs to generate simulated BOLD data from each voxel under different attention conditions. We then repeated the multivoxel stimulus reconstruction analysis on this simulated data to model population results for the second layer of the model. This approach allowed us to perform virtual experiments to test how changes in the first layer impacted the second layer. That is, we manipulated which vRF parameters changed with attention (first layer) and observed the resulting changes in the population-based stimulus reconstructions (second layer). For example, we could test whether an overall increase in vRF response gain with attention would be necessary or sufficient to reproduce the amplitude increases observed in the empirical stimulus reconstructions reported in Fig 5. These virtual experiments also allowed us to compare the relative impact of one type of response modulation (e.g. size changes) with other types of response modulations (e.g. position shifts).

Since the population-level stimulus reconstructions require many voxels from each subject to produce stable and reliable results, we combined the data across several regions in each individual subject before estimating the IEM (see **Materials and Methods** for a longer discussion). This yielded one occipital region that combined data from areas V1, V2, V3 and V4, and one posterior parietal region that combined data from V3A/B and IPS0. We repeated the IEM analysis described in the previous section on these larger regions, and found that the pattern of attentional modulations observed earlier was consistent in the large ROIs (Figure 5). Next, to verify whether we could perform the layered IEM using a reduced number of voxels, we re-ran the IEM analysis but only used the data from voxels with well-fit vRFs. The reduced dataset with fewer voxels reproduced the main pattern of results we observed in the previous section. In particular, covert attention led to an increase in the amplitude of reconstructions near the locus of attention (Fig 6b, black vs. gray bars).

We then investigated the contribution of each vRF parameter to the population-level stimulus reconstructions, in a comparison akin to the spatial discriminability analysis in Fig 3. A model that only simulated the observed vRF amplitude or vRF size modulations did not predict the observed increase in reconstruction amplitude near the attention target (Fig 6b, red lines). However, a layered model that only simulated vRF position changes did predict a large increase in reconstruction amplitude near the attention target in the parietal ROI (Fig 6b, blue line on right). This is consistent with the effects observed in the full dataset (Fig 5, Table 3), where we only observed an interaction of stimulus distance and attention in the parietal ROI.

To more formally quantify each manipulation of the layered IEM, we calculated an error metric to describe how well each model reproduced the attentional modulations in the empirical data (using the root mean square error, or RMSE). We compared each model’s RMSE to a baseline model, which did not simulate any vRF attentional modulations (far left in Fig 6c). This null baseline should have the highest error, and any good models should decrease the RMSE between the simulated data and the empirical data. Conversely, a model with higher RMSE is worse at accounting for the empirical data compared to the null model. In both the occipital and parietal ROIs, adding vRF position shifts to the layered model decreased RMSE, while abolishing position shifts generally increased the model error (Fig 6c). These data are consistent with the results from the spatial discriminability analysis. Altogether, they suggest that shifts in vRF position have the largest impact on population-level representations, while changes in vRF size or gain play smaller roles in changing the fidelity of the population code.

## DISCUSSION

By simultaneously measuring the response properties of both single voxels and populations of voxels within retinotopic areas of visual cortex, we could link attentional modulations of spatial encoding properties across scales. Our data provide an initial account of how different types of RF modulations improve the quality of population codes for visual space. First, we show how vRF attentional modulations depended on the distance between the vRF’s preferred position and the static attention target (Fig 2). We then found that shifts in the preferred position of vRFs near the attended target increased the spatial discrimination capacity of a population of voxels (Fig 3), as well as the amplitude of stimulus reconstructions based on response patterns across all voxels in a ROI (Fig 5, 6).

### Attentional modulations of spatial RFs

We provide new data on how vRF responses are modulated around a covertly attended static target (Sprague and Serences, 2013; de Haas et al., 2014; Klein et al., 2014; Kay et al., 2015; Sheremata and Silver, 2015). Like prior macaque studies, we find that vRF position shifts depend on the vRF’s distance from the attended target (Connor et al., 1996, 1997). However, we also found that the pattern of attentional modulations differs across the visual hierarchy. In V4, V3A/B, and IPS0 voxels shift towards the attended target, while in earlier areas, vRFs near the attended target are slightly repelled from it (Fig 2e). We also found distinct patterns of size modulations: vRF size increased near the attention target in early visual areas, but decreased in parietal areas V3A/B and IPS0. Comparison to the existing literature suggests that patterns of RF size modulations likely depend on the nature of the spatial attention task. In fMRI tasks where subjects attended to the mapping stimulus, rather than a static position, researchers report average vRF size increases with attention (Sprague and Serences, 2013; Kay et al., 2015; Sheremata and Silver, 2015). RFs in macaque area MT shrink when measured with a mapping probe smaller than the stimulus, but increase in size when macaques track the mapping probes as they move across the screen (Womelsdorf et al., 2006, 2008; Anton-Erxleben et al., 2009; Niebergall et al., 2011). This may be because the RFs shift position to track the probe, causing an apparent increase in overall size. Lastly, manipulating perceptual load at fixation also increases vRF size in human visual cortex (de Haas et al., 2014). Taken together, these observations demonstrate that the pattern of RF response modulations depends both on task demands and on the spatial relationship between the attended target and the encoding unit’s RF.

We note that while the similarity between attentional modulations of single cell RFs and single voxel RFs is compelling, their properties are derived from different input signals, and are not interchangeable. fMRI voxels in retinotopically organized regions of visual cortex sample from a broad array of neurons with roughly the same spatial tuning preferences, so a position shift in a vRF could either be driven by a change in the preferred position of single neurons, or by a change in the gain profile across neurons tuned to slightly different locations in the visual field. Similarly, single neuron RFs receive input from smaller RFs in earlier visual areas, and a position shift could arise from either mechanism described above (McAdams and Maunsell, 1999; Baruch and Yeshurun, 2014; Dhruv and Carandini, 2014). Because of this inherent ambiguity when measuring the encoding properties of a locally tuned unit, it is useful to compare them with attentional modulations of population-level representations.

### Attention boosts the spatial encoding fidelity of a population

We first measured the overall capacity of a population of voxels to make fine spatial discriminations in a region of space. We found that attention increased spatial discriminability near the attended target, relative to the ignored target. We then performed virtual experiments on the vRFs contributing to the population to determine how they affected the spatial discriminability metric. We report that vRF position shifts increased spatial discriminability significantly more than vRF size changes or gain changes (Fig 3).

Since the spatial discriminability metric (Fig 3) is only informative about a local portion of space, we performed a second population analysis to reconstruct an image of the entire visual field on each trial using a multivariate IEM. Attention increased the amplitude of stimulus reconstructions near the attention target, indicating an increase in representational fidelity that accompanied the increase in spatial discriminability. In addition, a layered spatial encoding model revealed that shifts in vRF position could account for these attentional enhancements in the population-level stimulus reconstructions, but changes in vRF size could not. Altogether, our data demonstrate that shifts in position of many RFs may be a dominant way that single encoding units alter the properties of a population spatial code.

Although population-level information increased the most with changes in vRF position, we reiterate that these position changes could arise from spatially-specific patterns of gain modulations in input RFs. If this is true, it is possible that gain modulations with attention may exert their largest effects on the downstream population, where these patterns of gain changes become apparent shifts in vRF position. However, this remains an open question for future work to address.

Our findings also underscore the fact that changes in the spatial encoding properties of single units do not directly translate into analogous changes in the encoding properties of a population of those same units. For example, an overall change in vRF size does not necessarily change the size of the population-level representation (Sprague and Serences, 2013; Kay et al., 2015). Although we found that single units shifted their preferred position towards the attended target, population-level representations did not generally shift with attention. When the population code did shift its encoded position, we found that it was typically representations of the ignored stimulus that shifted farther from the true stimulus location (Fig 5), consistent with more error-prone representations of irrelevant stimuli. These types of differences further emphasize the need to understand the effects of cognitive state on population codes for the entire visual scene, rather than focusing solely on single units.

Lastly, we note that our population-level data do not address the open question of whether RF attentional modulations have perceptual consequences, since it is not clear how the spatial encoding models measured here are linked to visual perception and behavior (Koenderink, 1990; Rose, 1999; Anton-Erxleben and Carrasco, 2013; Klein et al., 2016). Further investigation into these topics should include task manipulations to investigate how attentional modulations of both vRFs and population-level metrics track psychophysical performance.

### Tuning shifts and labeled lines

Historically, shifts in the tuning of a RF have not been considered one of the main mechanisms by which attention modulates population-level information, although recent reports suggest that this view is being reconsidered (David et al., 2008; Anton-Erxleben and Carrasco, 2013). This may be due to ‘labeled-line’ theories of visual information processing, which posit that a single neuron has a consistent feature label which downstream neurons rely on to perform computations and transmit stable information (Barlow, 1972; Doetsch, 2000; David et al., 2008). When a spatial RF shifts position as a function of cognitive state (e.g., attention), that single neuron’s feature label is no longer consistent. Without an accompanying shift in the downstream neurons receiving the changing feature label, such a change could disrupt the stability of the population code. However, our results suggest that population-level spatial representations remain relatively stable – and are even enhanced – when the tuning of the underlying vRFs shift in position, size, and gain.

An alternate proposal to a labeled line code relies on the joint information encoded across a population of cells (Erickson, 1982; Doetsch, 2000). This may occur at several scales–for example, V2 could use the pattern of information from V1 inputs to form a visual representation. This idea is more akin to an encoder-decoder model in which the downstream decoder does not need information about the altered representations in each of the encoder units, but instead relies on a population readout rule (Seriès et al., 2009). The population readout rule could incorporate knowledge about the ‘labels’ of the encoder units, but could perform equally well by relying on relative changes in the pattern across units to resolve the information encoded in the population. However, further exploration of population readout rules in visual cortex are needed to test this hypothesis.

## Conclusions

The spatial encoding properties of the visual system can be measured and modeled at many different spatial scales. Here, we report how these properties change with attention for single voxels and for a group of voxels in each ROI. Notably, single vRF modulations do not propagate directly to analogous changes in large-scale codes. Instead, we observed that attentional modulations of vRF position play a dominant role in modulating the amplitude of population-level representations. Future research is needed to resolve how shifts in RF labels are generated, how information is read out from a population, and how these multi-scale attentional modulations affect visual perception and behavior.

## Acknowledgements

Many thanks to the lab and particularly to Rosanne Rademaker and Edward Vul for comments on analyses and on the manuscript. This work was supported by National Science Foundation Graduate Research Fellowships to V. A. V. and T. C. S., and a grant from the National Eye Institute (R01-EY025872) and a Scholar Award from the James S. McDonnell Foundation to J. T. S.

## SUPPLEMENTAL METHODS

### Raw data and analysis code

All the data and analysis code needed to reproduce the analyses in this text are available in an Open Science Framework repository at https://osf.io/s9vqv/.

Population analysis (3): Layered spatial encoding model in smaller retinotopic ROIs

In the main text, we merge several retinotopically-defined ROIs to form a large occipital and parietal region before estimating the layered encoding model. When we attempted to estimate a layered IEM for the smaller ROIs, we were forced to exclude several participants because they did not have enough voxels in that region to calculate a stable population-level estimate of the spatial information in the mapped region. That is, the weight matrix estimated in the training portion of the IEM was poorly conditioned, or not full rank (Eq. 4). This resulted in the exclusion of 16 out of 42 possible participant-ROI pairs: V1 (AL); V3 (AL); V3A/B (AL, AP, AR, AU); V4 (AA, AL, AR, AU); IPS0 (AA, AL, AP, AR, AT, AU).

Note that the chosen level of noise did not qualitatively impact the results. For example, rather than just adding Gaussian noise, we also created a noise model that followed the covariance structure between all voxels in each ROI. To estimate the covariance matrix, we computed the residuals between the true trial-wise beta weights and the predicted trial-wise beta weights for each voxel based on its vRF model. We then calculated the pairwise covariance between the residuals for each set of voxels. Last, we added noise that followed this covariance structure to each voxel’s channel weights and simulated BOLD response. This noise was scaled to be the same as the noise level that most accurately captured the real reconstruction data (i.e., mean noise is 0.5 standard units). The pattern of results between each of the model manipulations remained the same, so those results are not discussed here.

#### SUPPLEMENTAL TABLES AND FIGURES

**Figure S1.**
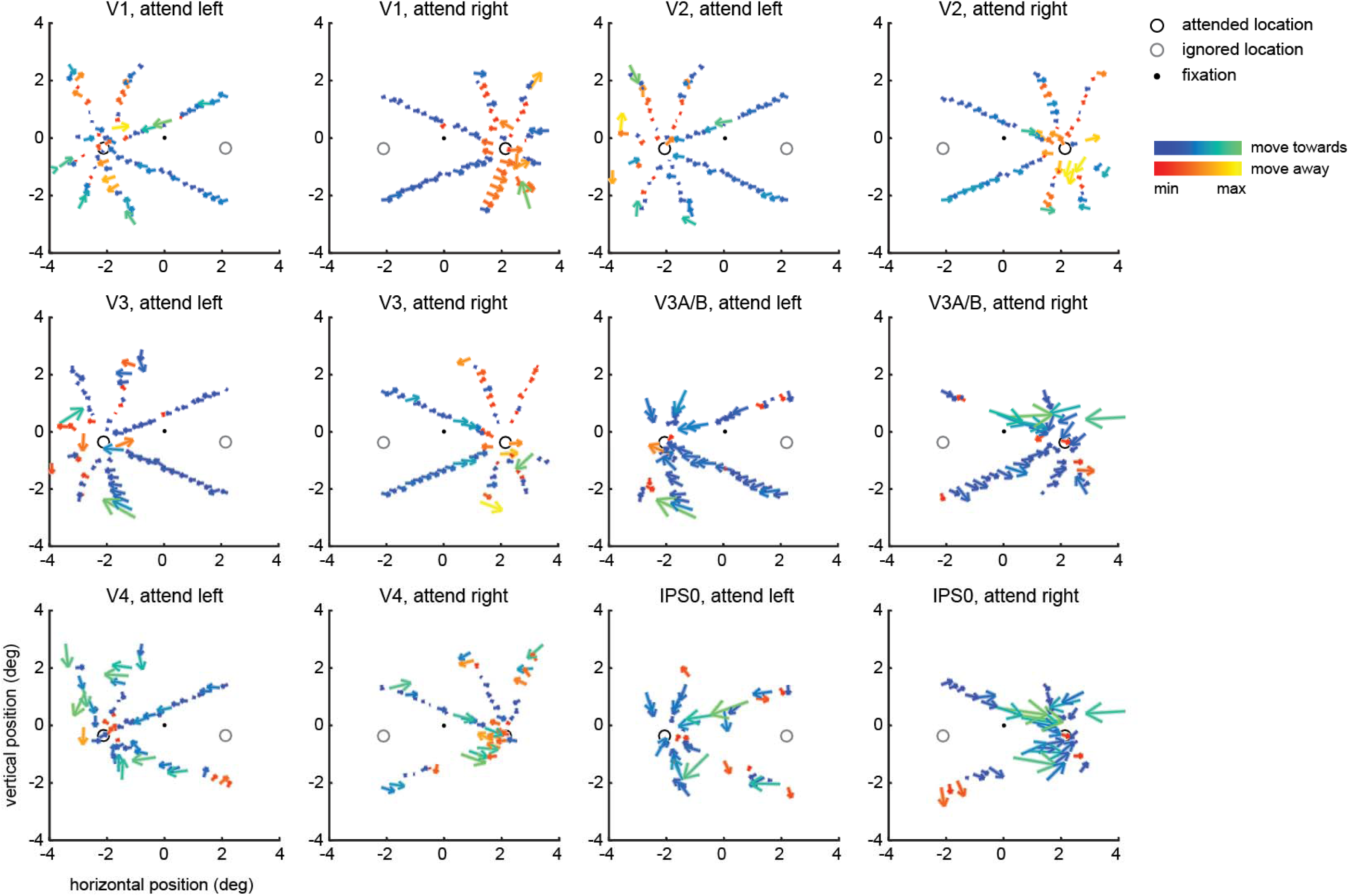
Preferred position changes of vRFs from each mapped visual area for both attention conditions. Like Figure 2C, these plots show participant averages. The majority of vRFs shift toward the attended location.

**Figure S2.**
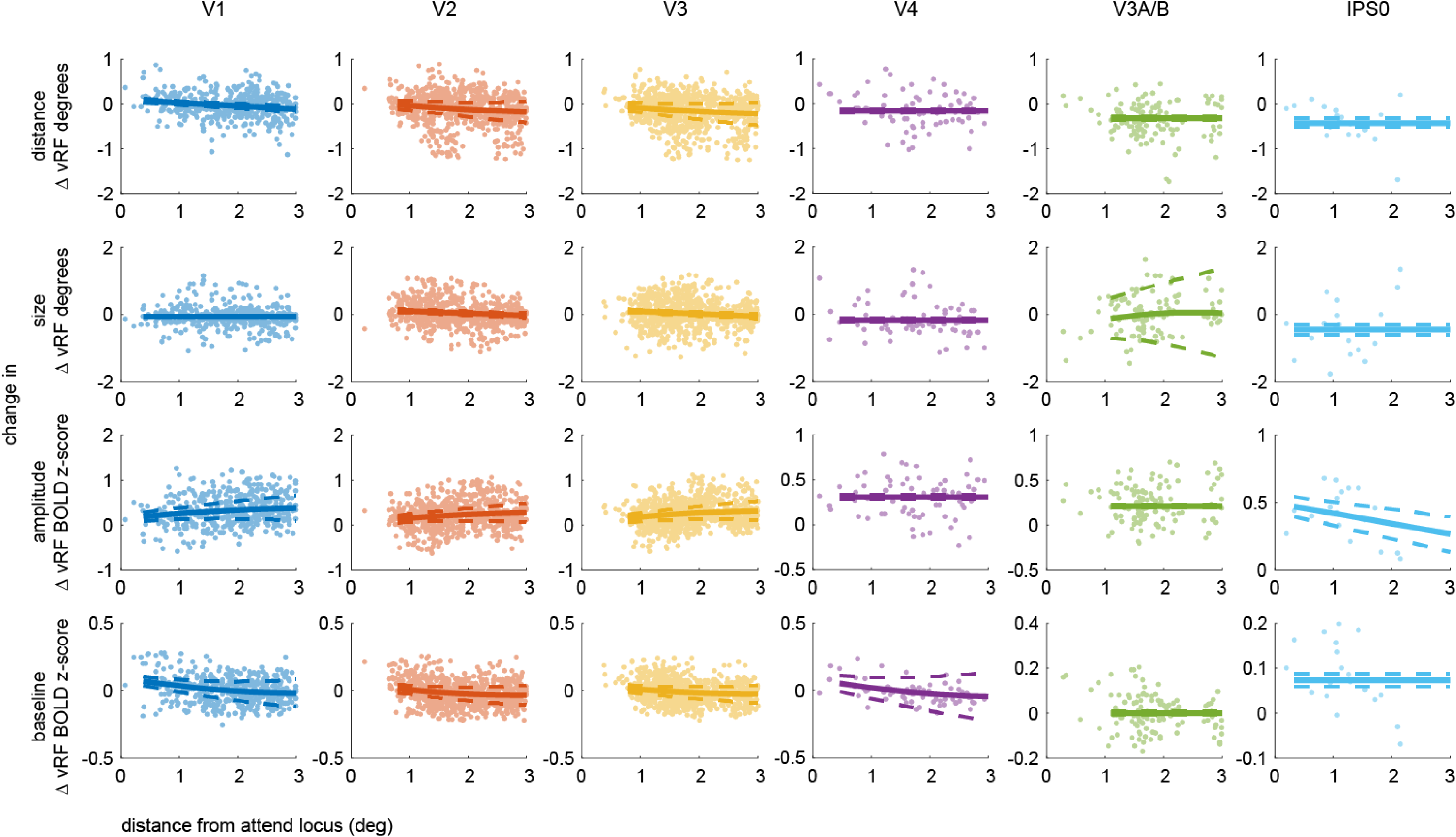
Related to Table 2. vRF attentional modulations as a function of distance from the attend location. All vRFs across participants are plotted here, overlaid with the best bootstrapped polynomial fit for every VOI-parameter pair.

**Figure S3.**
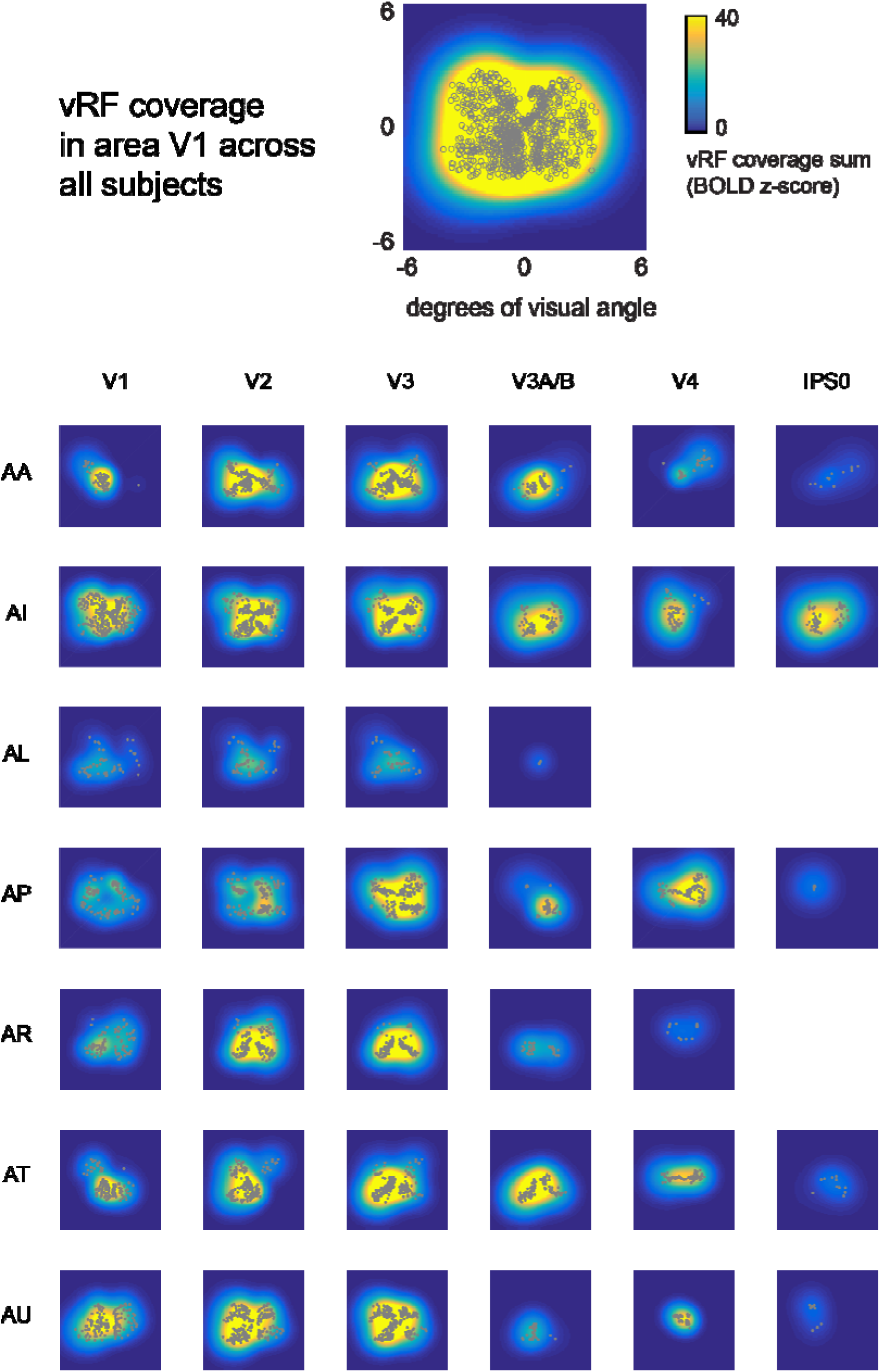
A plot of the visual field coverage of all ROIs and all participants for vRFs mapped during th eattend fixation runs. Top map shows the combined coverage across all participants in area V1. All image s are plotted on the same colorscale and only account for fit centers and sizes (e.g., no scaling by fit amplitude and baseline). Empty cells indicate that no voxels from that participant-ROI pair survived the voxel threshol ding procedure.

**Figure S4.**
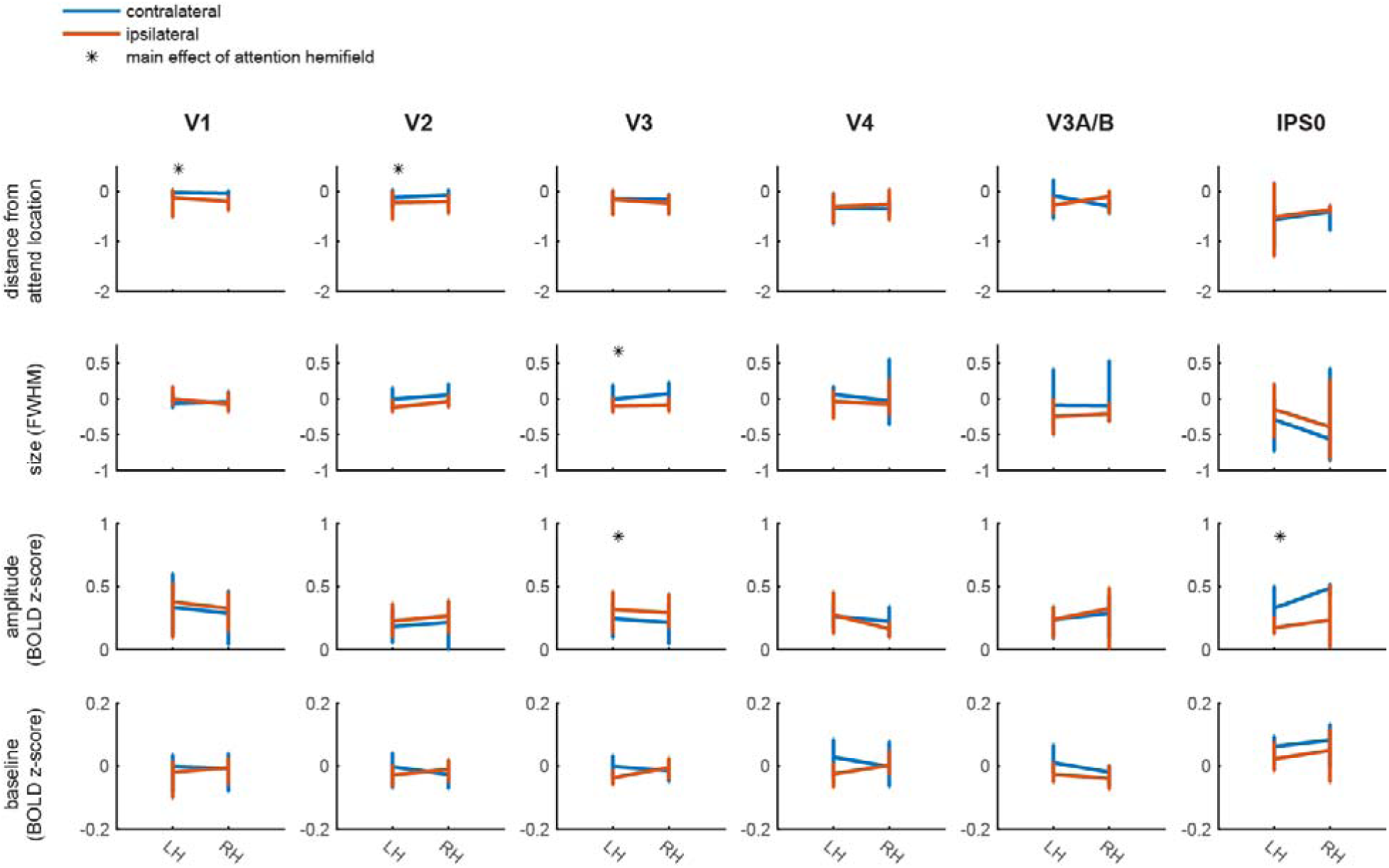
vRF attentional modulations by visual hemifield and voxel hemisphere. An asterisk indicates a maineffect of attended visual hemifield (see Table S3 for p-values). Areas with a significant difference in contralateral vs. ipsilateral difference scores are the same regions which have large slopes in **Fig S2**. This simply reflects the fact that voxels near the attended location are always contralateral (e.g., voxels in the RH will code for locations near the attention target in the left, or contralateral, hemifield). There was no main effect of voxel hemisphere or an interaction of the two factors.

**Figure S5.**
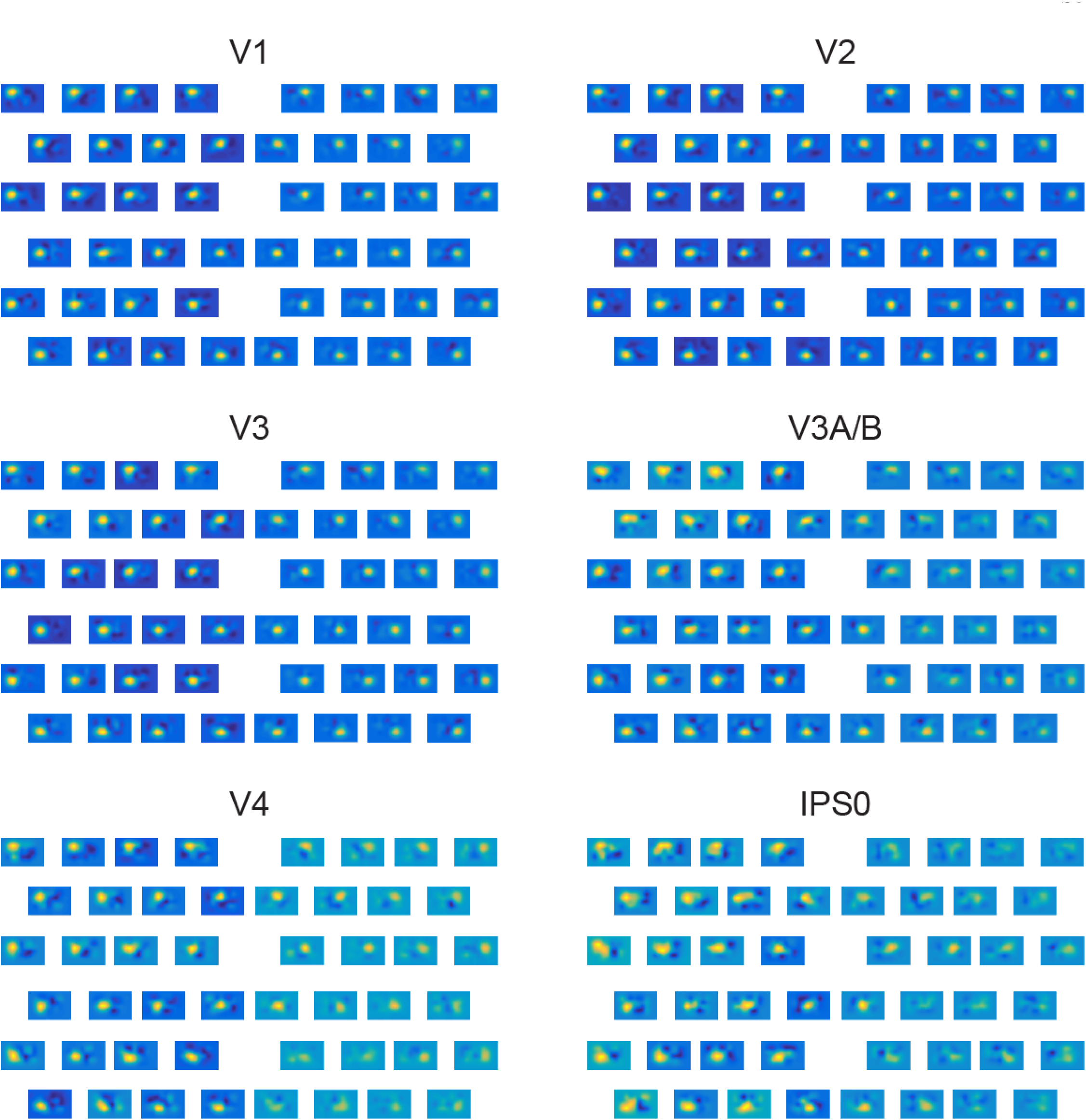
Stimulus reconstructions for each ROI, averaged across participants, and like positions across condition. Colorscale is constant across all 48 stimulus positions within an ROI. (Reconstructions for the attend right condition were flipped and averaged with the attend left condition.) The left hemifield is attended and the right hemifield is ignored. Stimuli that fall along the horizontal midline are excluded here.

**Figure S6.**
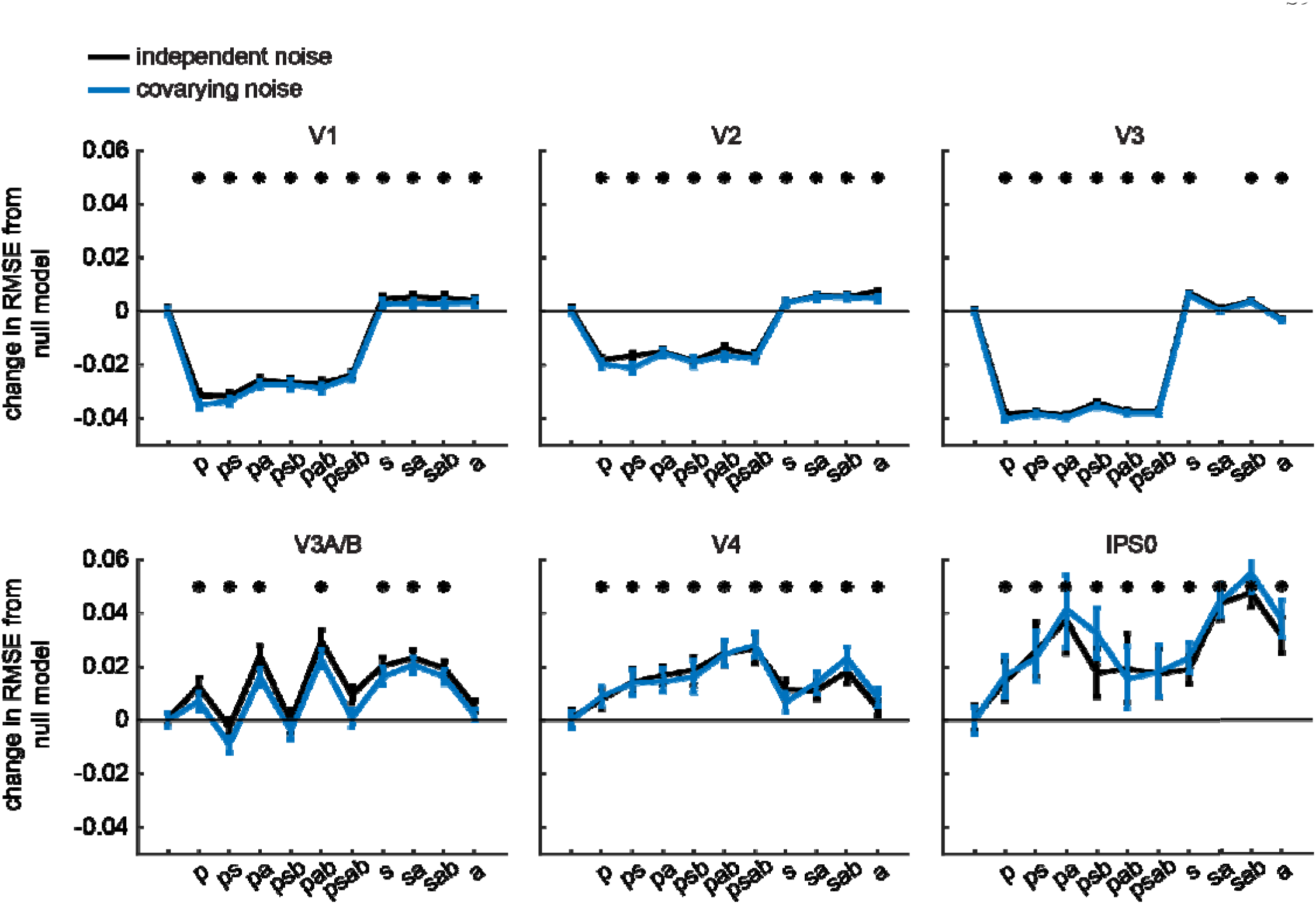
The level of noise added to the generated voxel responses was systematically manipulated (see **Supplemental Methods**). We then tested whether the noise parameters affected which model best explained theattentional modulations observed in the stimulus reconstructions. Shown on the x axis is which of the 4 vRF parameters was allowed to vary between attention conditions. The pattern of results was the same across models which used independent noise in the BOLD data simulation or noise which scaled with the voxelwise covariance matrix.

**Table S1.** Mean parameters (and 95% CIs) fit to vRF size data as a function of visual eccentricity

|  | Baseline | Slope |
| --- | --- | --- |
| V1 | .49 [.46, .53] | 1.34 [1.30, 1.39] |
| V2 | .41 [.36, .47] | 1.60 [1.53, 1.69] |
| V3 | .43 [.39, .47] | 1.86 [1.81, 1.92] |
| V4 | .76 [.67, .85] | 1.84 [1.72, 1.95] |
| V3A/B | .55 [.46, .63] | 2.20 [2.10, 2.29] |
| IPS0 | .92 [.69, 1.18] | 2.22 [1.78, 2.58] |

**Table S2.**
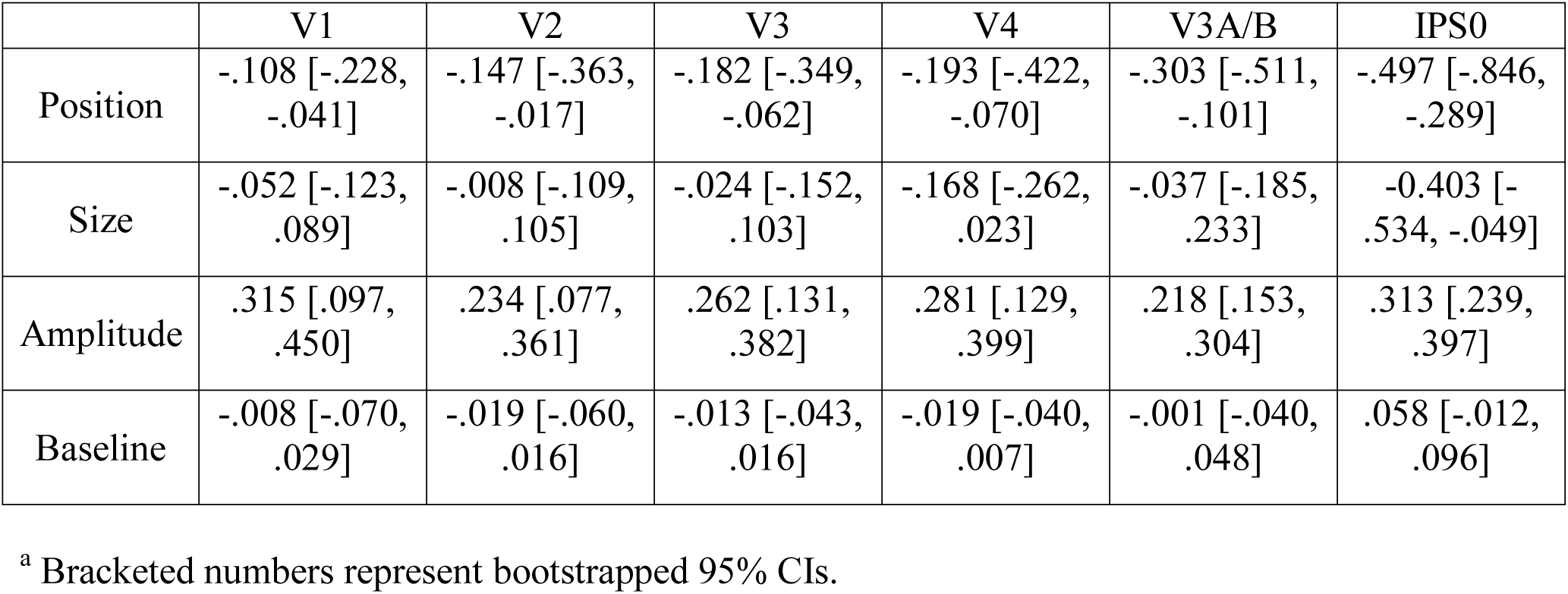
Overall changes in vRF parameters with attention (attend left or right–attend fixation)

**Table S3.** Exact p-values for the permuted tests of vRF modulations by voxel hemisphere and attention
hemifield, associated with Figure S4.

|  | V1 | V2 | V3 | V4 | V3A/B | IPS0 |
| --- | --- | --- | --- | --- | --- | --- |
| Main effect of attention hemifield |  |  |  |  |  |  |
| Position | **.007** | **.009** | .397 | .491 | .548 | .824 |
| Size | .884 | .081 | **.002** | .145 | .505 | .056 |
| Amplitude | .055 | .093 | **<.001** | .274 | .330 | **.005** |
| Baseline | .727 | .642 | .800 | .118 | .397 | .014 |
| Main effect of voxel hemisphere |  |  |  |  |  |  |
| Position | .435 | .702 | .572 | .756 | .770 | .648 |
| Size | .586 | .023 | .320 | .980 | .640 | .569 |
| Amplitude | .513 | .534 | .672 | .212 | .263 | .187 |
| Baseline | .715 | .892 | .385 | .129 | .813 | .187 |
| Interaction of hemisphere & hemifield |  |  |  |  |  |  |
| Position | .602 | .752 | .472 | .186 | .770 | .684 |
| Size | .103 | .726 | .476 | .654 | .874 | .571 |
| Amplitude | .710 | .965 | .887 | .367 | .699 | .275 |
| Baseline | .125 | .073 | .049 | .685 | .056 | .618 |

<sup>a</sup> **bold** numbers indicate that the p-value passed FDR-correction ( $q = .05$ , corrected across ROIs and comparisons within each parameter).

**Table S4.**
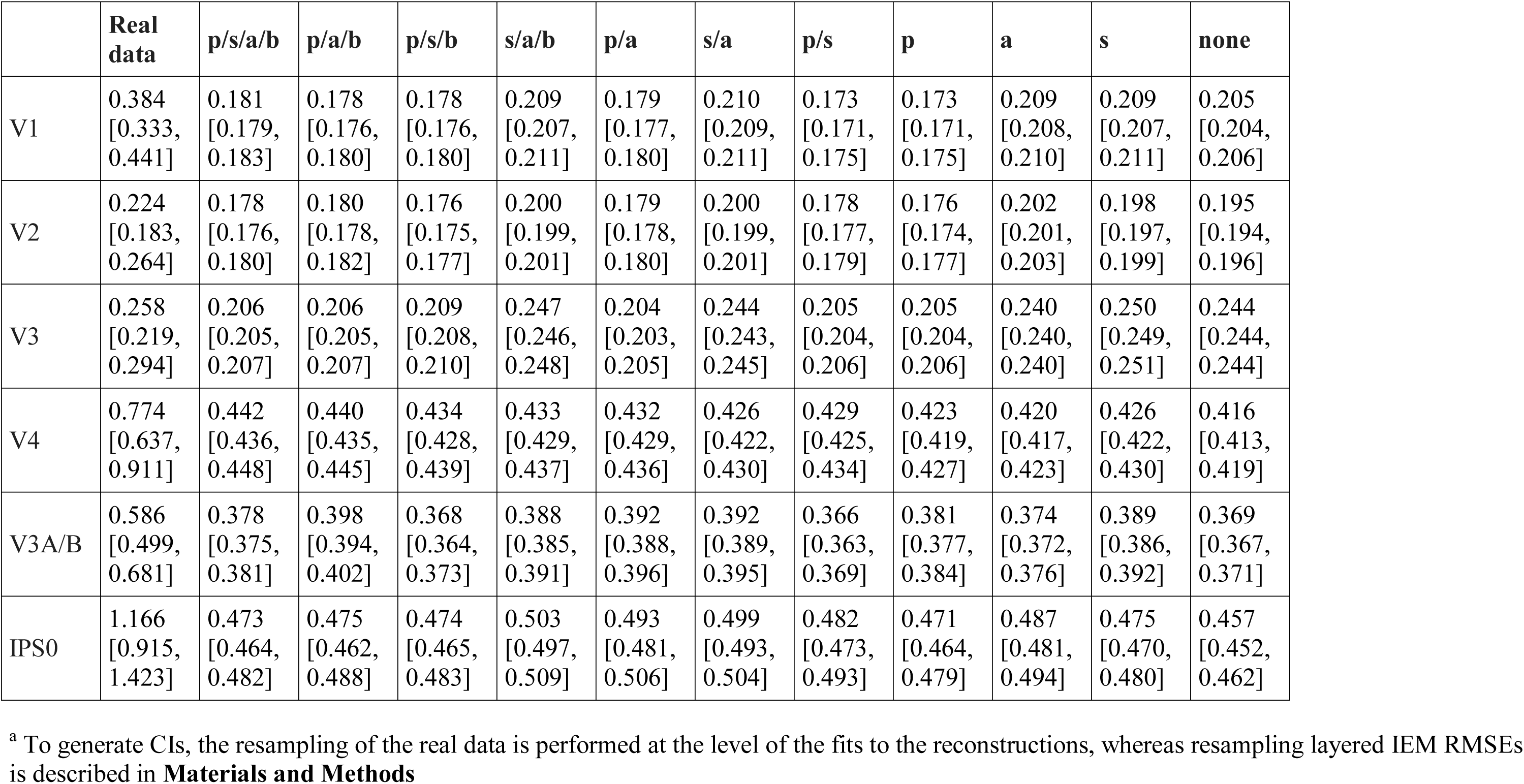
Model RMSE (and 95% CIs) between reconstructions from the reduced dataset (only using voxels with RFs) or from different versions of the layered IEM using the same voxels in the smaller retinotopic regions (compare to Figure S6).

